# Forecasting sockeye salmon migration timing and mortality under thermal stress

**DOI:** 10.64898/2026.11.20.765466

**Authors:** Nicholas A. W. Brown, Erika Eliason, Patrick L. Thompson

## Abstract

1. Pacific salmon hold high ecological, cultural, and economical importance, but climate change is constraining their survival and reproductive success. Warming rivers are one of the fore-most threats, with many populations increasingly facing temperatures approaching or exceeding their physiological limits. Understanding and predicting these negative impacts can inform mitigation or offsetting measures.
2. We used a 12-year dataset of 13 801 individually-tagged fish to develop a detailed model of Okanagan sockeye salmon migration based on functional relationships with river temperature and flow. Okanagan sockeye are the largest of the last two self-sustaining sockeye populations in the Columbia River basin. Their abundance increased dramatically over the last two decades amid significant rehabilitation efforts. However, Okanagan sockeye migrate into one of the species’ warmest inhabited regions and are highly vulnerable to climate change.
3. Our model revealed temperature thresholds associated with migration cessation and mortality near 21°C and 25°C, respectively. These temperatures are common during the migration period in the Okanogan River, which sockeye must ascend to reach the spawning grounds.
4. We used the estimated functional relationships to parameterize forward simulations of Okanagan sockeye migration from 2006–2026, spanning all years where water and fish passage data were available. These simulations accurately predicted migration timing and survival through the Okanogan River, providing a forecasting tool for Canadian fishery managers to regulate harvest in Osoyoos Lake.
5. The simulations also showed elevated mortality rates during warm years and revealed a potential ecological trap where brief periods of river cooling trigger large influxes of salmon migrants who may subsequently be exposed to extreme temperatures and experience high mortality rates.
6. Our work highlights the potential for functional models to provide detailed and accurate reconstructions and forecasts of Pacific salmon migration success, which managers will require as river temperatures increasingly exceed salmon physiological limits.

## Introduction

Climate change is driving global shifts in river temperatures (van Vliet et al. 2013), which pose acute threats to fish and other ectotherms whose physiology is controlled by ambient temperatures (Pörtner and Farrell 2008). Warming rivers are increasingly impacting Pacific salmon populations (*Oncorhynchus spp.*), particularly at southern latitudes where average temperatures are already near their upper thermal tolerance thresholds (Martins et al. 2011, Crozier et al. 2019). Exposure to supra-optimal temperatures inflicts metabolic, cardiorespiratory, immunocompetence, and reproductive costs on fish, hampering growth, reproductive success, and survival (Whitney et al. 2016). For Pacific salmon, these risks appear to be highest during upstream migration and spawning due to the physiological demands of swimming against high velocity flows (Eliason et al. 2011). Previous episodes of extreme river temperatures have been implicated in mass enroute mortality events (Farrell et al. 2008, von Biela et al. 2022), which pose serious complications for resource managers attempting to balance harvest against climate change constraints. Salmon fisheries are generally regulated using abundance forecasts with precautionary adjustments for subsequent natural mortality. However, these fixed adjustments may no longer be sufficient to achieve spawner abundance objectives under increasingly variable and episodic migration mortality (Holsman et al. 2019). Accurate forecasts of en-route mortality thus provide valuable information to managers facing difficult trade-off decisions (Macdonald et al. 2010, Hague et al. 2011).

Okanagan sockeye salmon (*O. nerka*) populations, located within the Columbia River Basin, occur at the southern periphery of the species latitudinal distribution and are presumed to be among the most vulnerable to climate-driven warming (Hyatt et al. 2003, Rand et al. 2012, Stiff and Thompson 2026). However, until recently, other anthropogenic pressures posed an even greater risk to Columbia River populations. Hydroelectric dam construction, irrigation diversion, pollution, invasive species introductions, and numerous other human impacts throughout the 1900s resulted in the extirpation of 5 upper Columbia River sockeye populations, critically endangered status for the Snake River population, and depressed abundances of the Okanagan and Wenatchee populations (Gustafson et al. 2007, Rand et al. 2012, Hyatt and Stockwell 2019). To address these issues, a suite of management and restoration efforts were implemented in the early 2000s that, together with favourable marine conditions, instigated an approximately four-fold increase in Okanagan sockeye abundance beginning around 2008 (Alexander et al. 2024, Bailey et al. 2025). As a result of this impressive recovery, the Okanagan population has recently supported expanded First Nation and recreational fisheries in Canada (Hyatt and Stockwell 2019). The present challenge for fishery managers is to ensure harvest rates and timing are appropriately adjusted to mitigate climate change impacts.

Currently, sockeye passage counts at Wells Dam in central Washington, USA, are the most proximate data informing the abundance of Okanagan sockeye returning to Canada (Ogden et al. 2025, Fisheries and Oceans Canada 2025), but their relevance to fishery managers is deteriorating. Wells Dam counts have historically correlated with subsequent spawning ground estimates, which justified their use in setting harvest levels (Hyatt and Rankin 1999). However, more frequent drought conditions and “thermal barrier” events (Mayer et al. 2024) arising from climate change are hampering sockeye survival and migration in the Okanogan River above Wells Dam, forcing managers to make heuristic adjustments attempting to compensate for delayed arrival timing and excess en-route mortality. This issue was highlighted during the significant drought of 2015, when anomalously warm winter temperatures and a low spring snowpack resulted in near-record high springtime Columbia River temperatures (Marlier et al. 2017, Crozier et al. 2020) and dramatic sockeye mortality rates. Although close to 450 000 Okanagan-bound sockeye returned to the mouth of the Columbia River in 2015 (the second highest return since the 1970’s at the time), only about 20 000 (5%) were subsequently estimated on the spawning grounds (Bailey et al. 2025, Ogden et al. 2025, Fryer et al. 2017). (The endangered Snake River sockeye population similarly suffered approximately 92% en-route mortality (Crozier et al. 2020)). While 2015 constitutes an extreme outlier in reference to historical conditions, it represents average expected conditions in the mid-21^st^ century (Marlier et al. 2017, Stiff and Thompson 2026). Fishery managers thus require accurate predictions of migration delays and natural mortality rates above Wells Dam to inform sustainable harvest levels and opening times moving forward.

Our objective was to support climate adaptation in Okanagan sockeye fishery management by developing an environmentally-conditioned forecast of the Okanagan sockeye arrival timing and total return abundance to Canada. We began by developing a Bayesian model linking migration outcomes to river temperatures and flows, which uncovered key thermal thresholds associated with migration cessation and mortality. We then used the estimated functional relationships from this model to reconstruct historical migration dynamics using simulations driven by river conditions. Finally, we compared the simulated and observed migration dynamics to evaluate potential forecasting performance.

## Methods

### Okanagan sockeye salmon upstream migration

Okanagan-bound sockeye salmon enter the mouth of the Columbia River in June and July. Their journey to the spawning grounds above Osoyoos Lake spans more than 1000 km with 1 km of elevation gain, and traverses 9 dams on the Columbia mainstem, 1 dam on the Okanogan River, and up to 2 dams on the Okanagan River (Figure 1). The average migration time is about one month, with a corresponding swimming speed of about 30 km per day (Hyatt et al. 2003). Swimming speeds are generally sustained around 40–50 km per day between Bonneville and Wells dams (located 234 km and 830 km upstream, respectively), but then become slower and much more variable beyond Wells Dam, reflecting holding and migration delays at the transition from the Columbia mainstem into the Okanogan River. Warm Okanogan River temperatures often cause sockeye to hold for extended periods in the Wells Dam reservoir, Lake Pateros, which serves as a thermal refuge (Hyatt et al. 2003, Fryer et al. 2017, Murauskas et al. 2021). After entering the Okanogan River, sockeye must travel 125 km and surmount Zosel Dam to reach Osoyoos Lake and the Canadian border (Figure 1). Sockeye are harvested at various locations along their migration route in the Columbia River, including significant recreational and indigenous fisheries in Lake Pateros. Canadian recreational and indigenous harvest occurs in Osoyoos Lake.

**Figure 1:**
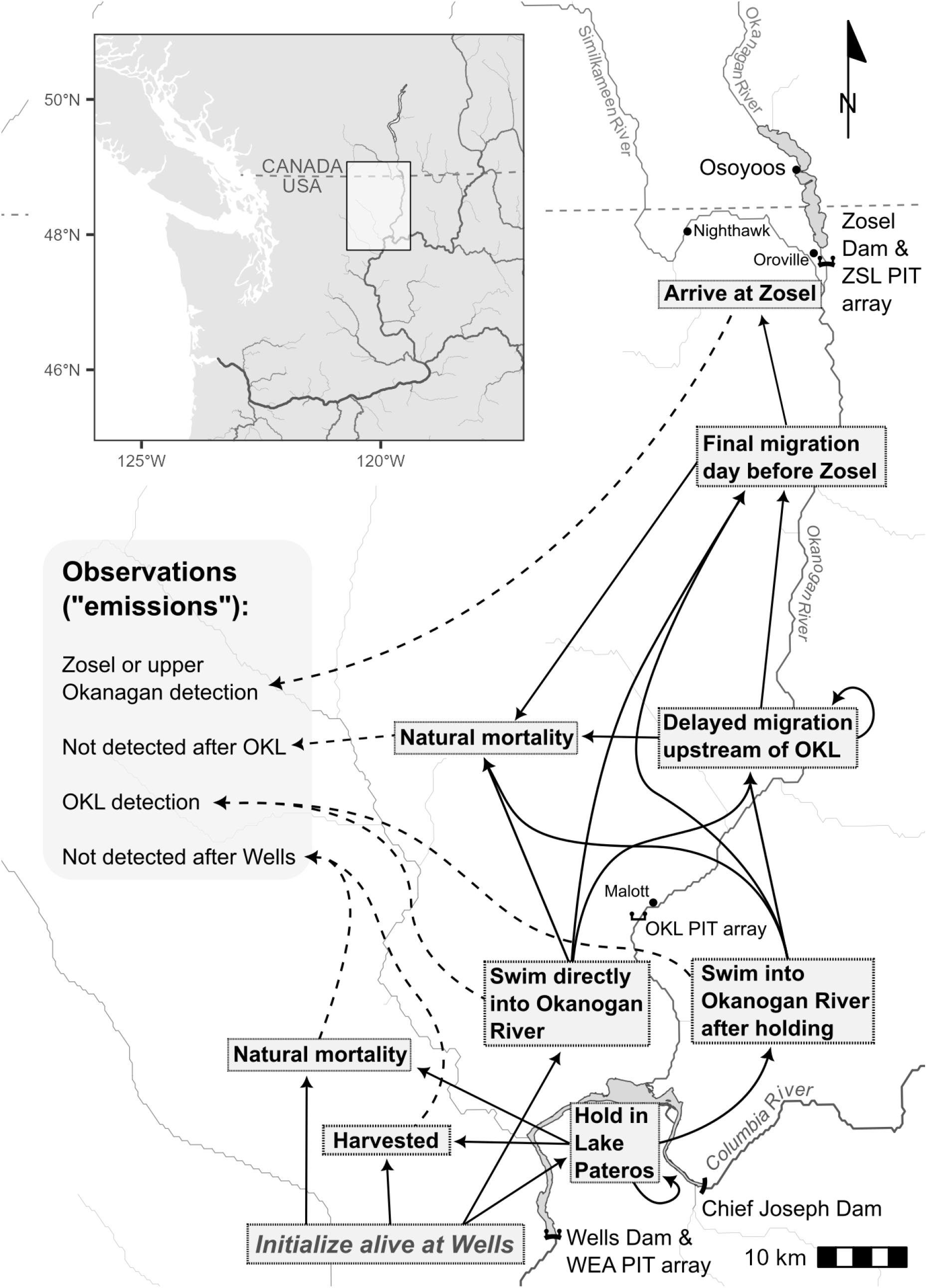
Map of the Okanogan River overlaid with HMM latent states and paths.

### Data sources

Sockeye migrants are counted daily at each of the Columbia mainstem dams. In addition, thousands of adult Sockeye are implanted with passive integrated transponder (PIT) tags each year in assessment programs primarily operating at Bonneville and Wells dams. Other assessment programs at Rock Island Dam and in Canada implant tens of thousands of juvenile sockeye with PIT tags prior to their seaward emigration; those who survive and return to the Columbia River then provide additional data as upstream migrants. We accessed adult passage count data from the University of Washington’s Columbia River DART portal (www.cbr.washington.edu/dart) and PIT tag data from the PTAGIS database (www.ptagis.org). PIT data were filtered to include only adult fish moving through the system in years where the Wells Dam, lower Okanogan River, and Zosel Dam arrays (“WEA,” “OKL,” and “ZSL,” respectively) were operating, which resulted in a final dataset of 13 801 tags across 2014–2025.

Hourly or daily river temperature and discharge data are collected at numerous stations along the Okanagan, Okanogan, and Columbia rivers. Most relevant to this study were data collected from the Okanogan River near Malott and Oroville, WA, from the Similkameen River near Nighthawk, WA, and from the Columbia River at Wells and Chief Joseph dams. We retrieved these data going back to 2006 using the Columbia River DART portal (link above) and the US Geological Survey’s ‘dataRetrieval’ R package (DeCicco et al. 2014). We imputed water temperature and discharge values on days with missing data to obtain complete time series for the key stations in our analysis—Malott and Wells Dam (see Supporting Information for data imputation details). Estimates of sockeye harvest between Wells Dam and Osoyoos Lake were provided by the Washington State Department of Fish and Wildlife. Recreational kept and released catch estimates were available on a monthly basis and other catch estimates were available on an annual basis.

### Migration dynamics model

#### Process model

Given that our primary objective was to characterize the impacts of the Okanogan River thermal barrier, we focused our model on migration above Wells Dam. We used Zosel Dam as the upstream boundary because it is the final observation point before sockeye reach Osoyoos Lake, where Canadian harvest occurs.

We assumed travel times and survival between Wells and Zosel dams are driven by four key factors: holding in Lake Pateros, migration delays in the Okanogan River (which could include slowed swimming or further holding in tributary thermal refugia; Goniea et al. 2006), human harvest, and natural mortality. Our approach was to model holding, migration delay, and natural mortality as functions of river temperature and flow using logistic regressions:

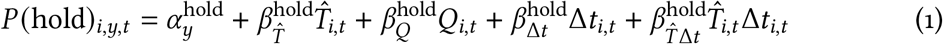

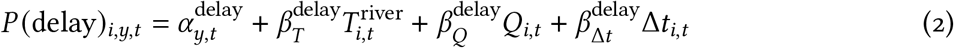

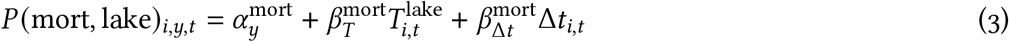

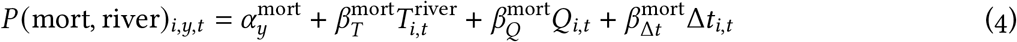

where *i*, *y*, and *t* subscripts denote terms that are indexed by fish, year, and day, respectively. *P* (hold), *P* (delay), *P* (mort, lake), and *P* (mort, river) are daily probabilities of holding in Lake Pateros, migration delay in the Okanogan River, dying in Lake Pateros, and dying in the Okanogan River, respectively. In each regression, *T*, *Q*, and Δ*t* represent daily mean water temperature (mean-centered at 18.8°C), daily mean log-flow (mean-centered at 3.67 log 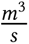), and elapsed days (scaled by 30 days and centered on day 5 for delay only), respectively; *T̂* is the rolling 4-day mean Okanogan River temperature (mean-centered at 19.9°C), reflecting both current conditions and the trend over recent days, which both prior research and our own exploratory analyses indicated are important for holding (Major and Mighell 1966). The *α_y_* terms are partially-pooled annual intercepts, *β* terms with single subscripts are slope coefficients for each covariate, and 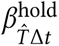 is an interaction coefficient between *T̂* and Δ*t*.

We modeled harvest as a daily latent competing-risk process affecting all vulnerable fish holding or migrating through Lake Pateros. Fishing encounters result in either kept harvest or release, so we partitioned latent annual fishing-encounter hazards, *F_y_*, between these two outcomes:

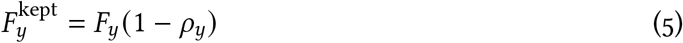

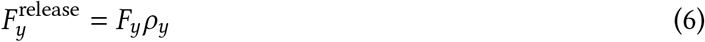

where *ρ_y_* is the latent annual release rate. Annual fishing-encounter hazards were converted to daily hazards, *H_y_*_,*t*_, using annually-normalized daily weights, *w_y_*_,*t*_, which convey the seasonal timings of reported kept harvest and releases:

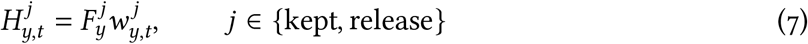

The daily probability of any fishing encounter was thus:

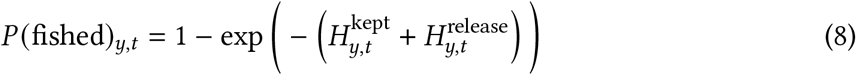

Conditional on a fishing encounter, daily competing-risk probabilities for kept harvest versus release were allocated in proportion to the component hazards:

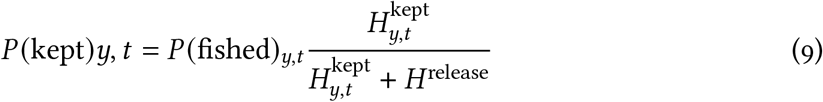

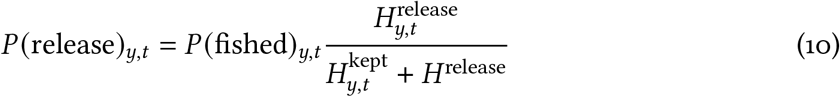

Released fish were vulnerable to fishing-related incidental mortality (FRIM; Patterson et al. 2017) with conditional probability *π*_FRIM_. Thus, the daily probabilities of FRIM following release were:

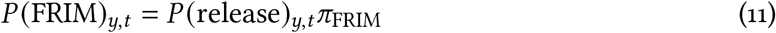

The challenge was connecting these equations to observed data. The PIT tags provide valuable information on travel times and survival of individual fish between detection arrays. However, the statuses of tagged fish are unknown for all days except when they are directly observed. We thus elected to represent the Wells-Zosel migration dynamics in a hidden Markov Model (HMM; McClintock et al. 2020), treating harvest, natural mortality, holding, and migration delays as latent unobserved states.

Latent states and pathways encoded in the HMM are depicted in Figure 1. Sockeye are exposed to fishing-encounter hazards (harvest or release) on initialization during their Wells passage day. Fish that survive either enter Lake Pateros holding or swim directly into the Okanogan River (taking up to 2 days post-Wells). Fishing-encounter hazards are applied each day fish are either holding in Lake Pateros or swimming toward the Okanogan River. Fish are assumed to enter the Okanogan River the day after holding ceases. Once in the river, fish either enter a delayed migration state, or immediately begin their final migration day before reaching Zosel Dam. State-specific mortality is applied daily after resolving fishing hazards. Zosel arrival and all mortality states (harvest, FRIM, and natural) are absorbing.

#### Observation model

PIT detections were used to distinguish latent paths. Detections at the OKL array were assumed to indicate same-day river-entry and detections at ZSL indicated same-day Zosel arrival (Figure 1). Disappearances between WEA–OKL or between OKL–ZSL were assumed to indicate mortality in Lake Pateros or in the Okanogan River, respectively (Figure 1). However, the ZSL and OKL arrays have poor detection efficiencies in some years (Fryer et al. 2017), so we included a Cormack-Jolly-Seber observation process to account for disappearances due to array bypass (Figure 1B; Whoriskey et al. 2019). To inform ZSL detection efficiencies, we pooled all 9 upstream Canadian arrays into a single array cluster ‘UOK’ (i.e. ‘upper Okanagan’) and retained the earliest detection date for tags with multiple observations. HMM latent states were thus linked to PIT detections accordingly:

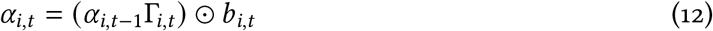

where *α_i_*_,*t*_ is a vector of forward state probabilities for tag *i* on day *t*, Γ is a daily transition matrix, and *b* represents observation compatibility and annual array-specific detection efficiencies. These calculations were performed in log space. Full transition equations and observation constraints for each latent pathway are provided in Tables S6 & S7.

#### Population-level harvest recursion

Because harvested PIT tags were not recovered within the model region, we used a simultaneous population-level forward recursion to inform annual fishing intensity, which ensured the estimated latent daily competing-risk probabilities included in the HMM—*P* (kept) and *P* (release)— were sufficiently constrained to reproduce the reported annual kept and released catch, within observation uncertainty bounds. In this recursion, daily Wells passage counts were treated as ‘cohorts’ and propagated through the same transitions as individual PIT-tagged fish. The recursion tracked daily expected vulnerable abundance, 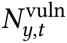, and accumulated expected kept and released catches over the year:

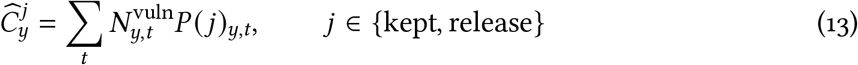

Observed annual kept harvest and releases, 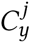, informed the latent fishing process through separate lognormal observation models:

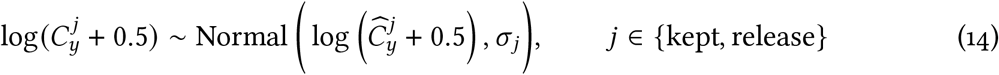

#### Hierarchical structure and priors

Regression intercepts for holding, mortality, and migration delay were partially-pooled across years, as were fishing-encounter hazards, release rates, array detection efficiencies, and mean ZSL-to-UOK survival.

Most priors were weakly informative. Regression intercept and slope terms for holding and mortality were constrained to plausible ranges based on thermal performance curves for Fraser sockeye populations (e.g. Eliason et al. 2011). Mean detection efficiency for the UOK PIT array cluster was centered on 95% to reflect generally strong confidence in the mainstem “OKC” array (E. McGrath, Okanagan Nation Alliance, pers. comm.) plus additional detection opportunities at the other 8 arrays. Prior distributions and bounds are listed in Tables S2, S3, & S4.

#### Sampling and diagnostics

We fit the HMM in a Bayesian estimation framework with R and Stan (Carpenter et al. 2017, R Core Team 2026). We sampled from 4 chains with 850 iterations each and discarded the first 600 iterations as a warm-up. Traceplots and ‘cmdstanr’ (Gabry et al. 2025) diagnostics indicated no divergent transitions. Bulk effective sample sizes averaged 1091 (range: 255–2527) and 80% of *R̂* were less than 1.01 (max: 1.037).

### Migration simulations

The HMM yielded functional relationships linking environmental variables to key migration outcomes, as well as daily fishing-encounter competing-risk probabilities. Our ultimate aim was to use these parameters to understand how environmental conditions and harvest cumulatively affect migration success of individual fish and of the population as a whole. We therefore translated the HMM process logic into two independent migration simulations in R—one for individual PIT tags and one for all fish enumerated at Wells Dam.

#### Forward-backward smoother on individual PIT tags

We used individual-level simulations to validate the process logic in the HMM. PIT-tagged fish, whose data were used to fit the HMM, were initialized and advanced through the latent states and transition probabilities using parameter values randomly sampled from the Stan model posterior. We then plotted the resulting data as time series showing each fish’s latent state progression in relation to Okanogan River temperatures (Figure 2). These simulations confirmed that the HMM infers plausible state histories for each fish.

**Figure 2:**
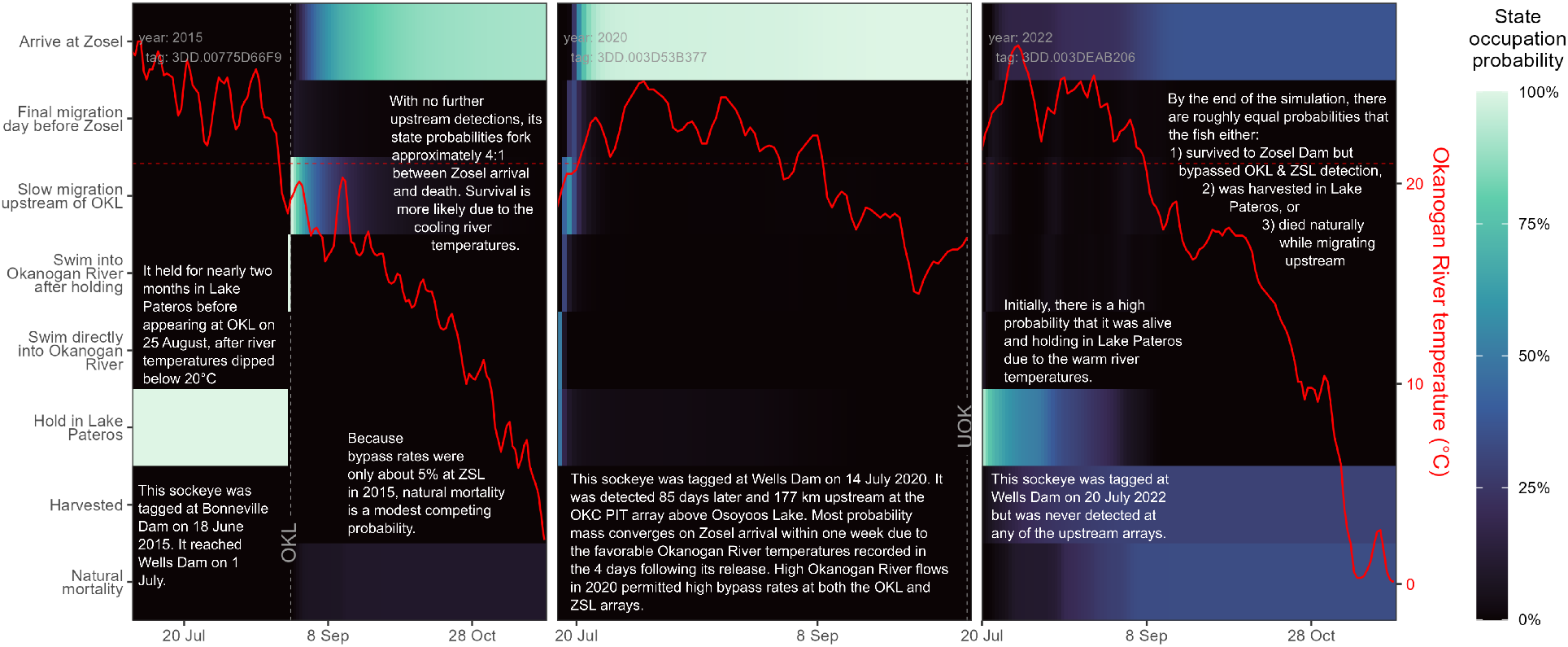
Simulated latent state probability time series for three sockeye PIT tags. Latent states are y-axis categories and dates are plotted along the x-axis. The red line shows daily Okanogan River temperatures and is plotted with respect to the right y-axis. The red dashed line corresponds to 21°C. Labeled, dashed vertical lines indicate observed detections at the stated array.

#### Population-level forward simulations: a return-to-Canada forecast

We used population-level simulations to propagate all sockeye counted past Wells dam through the HMM process logic to estimate the numbers of fish occupying each state across the migration season. We assumed the numbers flowing into the ‘Zosel arrivals’ state would be closely indicative of the numbers reaching Osoyoos Lake and the Canadian border, thus constituting a useful return-to-Canada prediction. In the simulations, each daily Wells passage cohort was initialized using the same competing risks and advanced through the same latent states and transition probabilities as in the HMM. We ran these simulations for all years where the required environmental, fish passage, and harvest data were available, 2006–present (2026).

Parameter values used in the simulations differed between years that were (2014–2025) versus were not (2006–2013 & 2026) directly informed by PIT data. For years with PIT data, hierarchical parameter values were randomly drawn from the matching year-specific posterior distributions. For years without PIT data, hierarchical parameter values were randomly drawn from across all year-specific posterior distributions. However, the annual fishing-encounter hazards could not be generalized across years because each is calibrated to annual reported kept catch and releases. Thus, for years without PIT data, we conducted year-specific calibrations to estimate *F_y_* and *ρ_y_* by running daily Wells counts through the forward simulation and searching for the root *F_y_* and *ρ_y_* values where simulated kept catch and releases were most similar to the reported totals.

Because the simulations were used for two different purposes—fishing-encounter hazard calibration and return-to-Canada forecasting—we required two different modes for applying state transitions: deterministic and stochastic. Under the deterministic mode, daily cohort abundances were multiplied smoothly across the daily state probabilities vector, providing necessary stability for *F_y_* and *ρ_y_* root-finding. Under the stochastic mode, a multinomial draw from the daily state probabilities vector was used to apportion all fish to a single state, producing more realistic noise in the daily predictions. The stochastic simulation outputs contain daily predictions for numbers of fish occupying each HMM latent state, with moralities apportioned among fishery harvest, FRIM, natural mortality in Lake Pateros, and natural mortality in the Okanogan River (Figure 5).

To assess forecast quality, we compared simulated data to observed daily sockeye counts through Zosel Dam, which were collected between 2006–2017. However, 2011–2013 and 2017 are considered unreliable due to recurrent equipment failures and unquantified bypass (Hyatt et al. 2020), so we included only the 2006–2010 and 2014–2016 data in our quality assessments. Within these years, some dates were missing observed data, so we also filtered these dates out from the simulated time series for consistency. We quantified forecast quality with continuous ranked probability scores (CRPS), a proper scoring rule that accounts for both the accuracy and precision of a predictive distribution (Gneiting and Raftery 2007). We calculated the CRPS of cumulative daily values across posterior draws passed to the simulation for each year and year–day combination. We divided the daily and annual CRPS values by the corresponding cumulative observed Zosel count values to convert them to percentage errors. To calculate smoothed expected forecast error values for each day of the sockeye migration season, we fit the daily percentage error values using a generalized additive mixed-effects model (GAMM) with a smoothed day effect and random intercepts for each year.

## Results

### Effects of river temperature and flow on migration dynamics

Sockeye behaviour and natural mortality are strongly linked to Okanogan River temperatures (Table 1). For fish that have just passed Wells Dam, daily holding probabilities of 50%, *P* (hold) = 50%, occur when the 4-day rolling mean Okanogan River temperature is approximately 20.5°C (95% credible interval [CI]: 20.2–20.8°C); *P* (hold) = 90% occurs around 22.4°C (22.2–22.7°C; Figure 3). However, these temperature thresholds decrease the longer fish have been holding. For example, after holding for 30 days, *P* (hold) = 50% and *P* (hold) = 90% occur around 19.5°C (19.1–19.8°C) and 21.9°C (21.6–22.2°C), respectively (Figure 3). Water temperatures also appear to modestly impede migration once sockeye enter the Okanogan River. Daily probabilities of continued migration delay, *P* (delay), increase from approximately 37% (27–50%) at 18°C to approximately 50% (37–62%) at 26°C (Figure 3). For natural mortality, *P* (mort) = 50% and *P* (mort) = 90% occur around 25.0°C (24.7–25.3°C) and 26.8°C (26.3–27.1°C), respectively (Figure 3).

**Figure 3:**
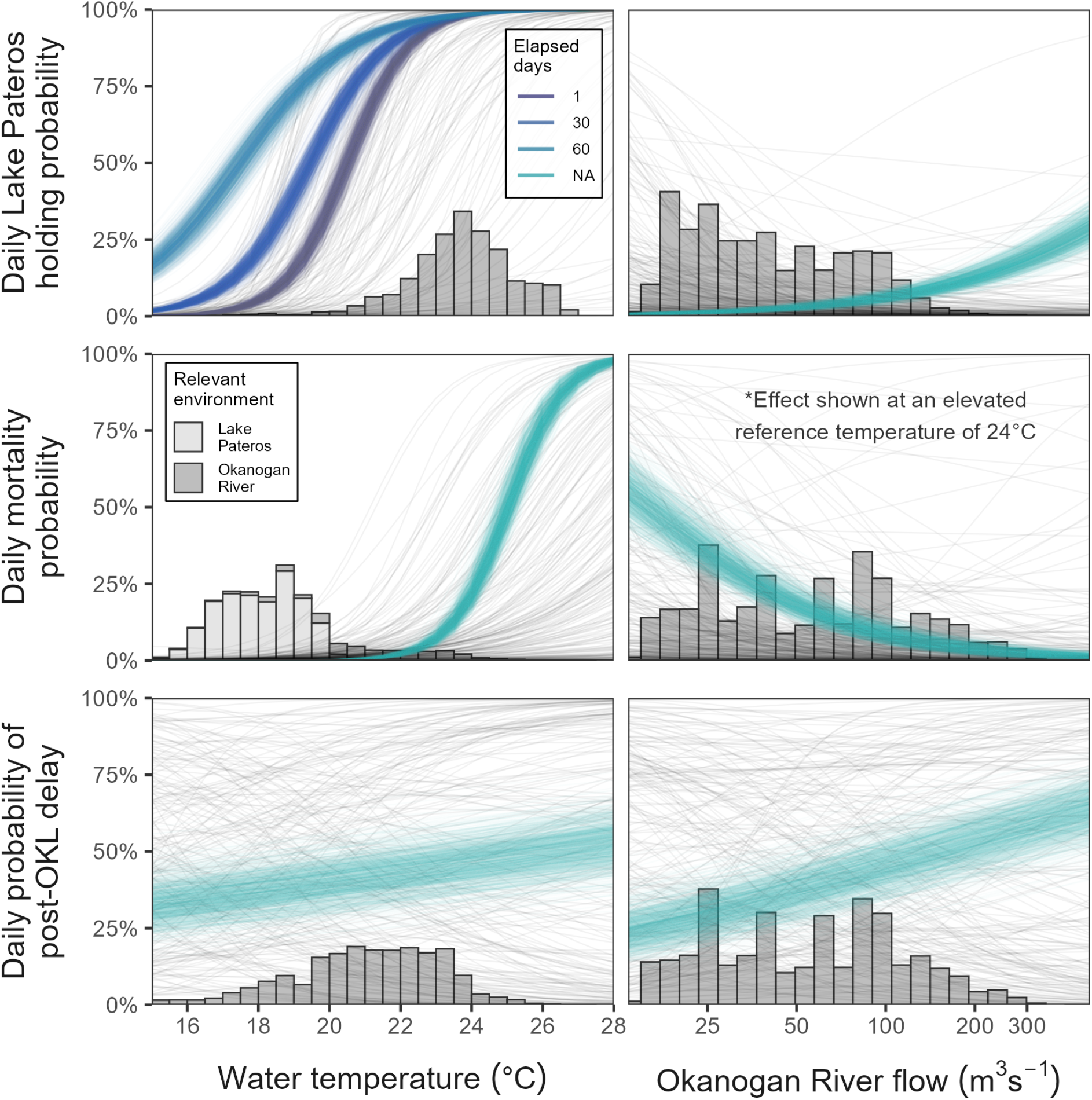
Okanagan sockeye daily holding, mortality, and in-river migration delay probabilities plotted as functions of water temperature, Okanogan River flows (log-transformed), and elapsed days post-Wells passage. Grey and coloured curves show prior and posterior draws, respectively. Daily holding probabilities versus water temperature are plotted at three different elapsed day values to show the mediating effect. Daily mortality hazards versus flow are plotted using a higher reference temperature of 24°C because mortality hazards are nearly 0% (regardless of flow) at the model intercept temperature of 19.9°C. Grey bars are density histograms showing posterior expected distributions of environmental conditions encountered while fish were at risk of the corresponding daily process.

**Table 1:**
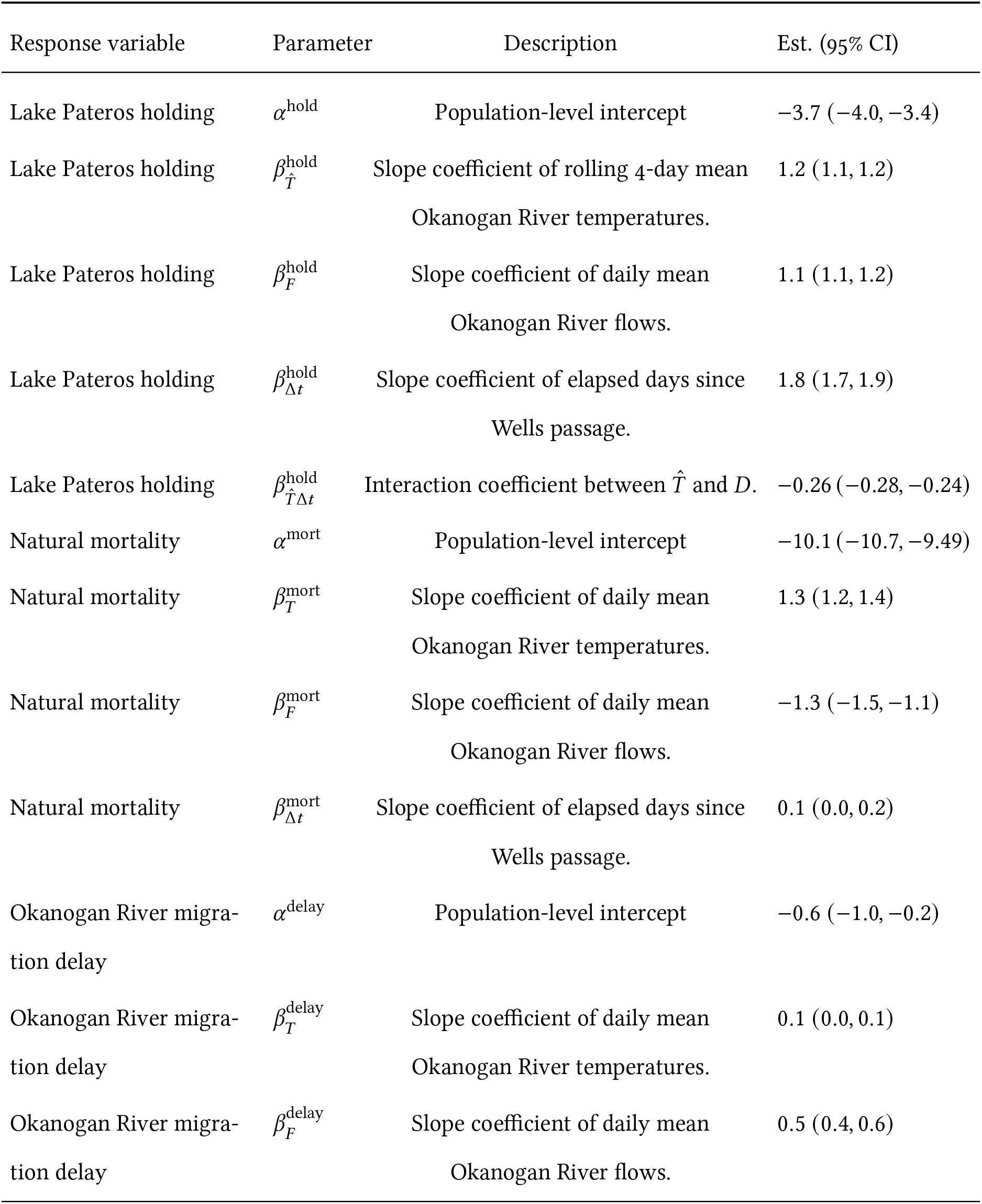
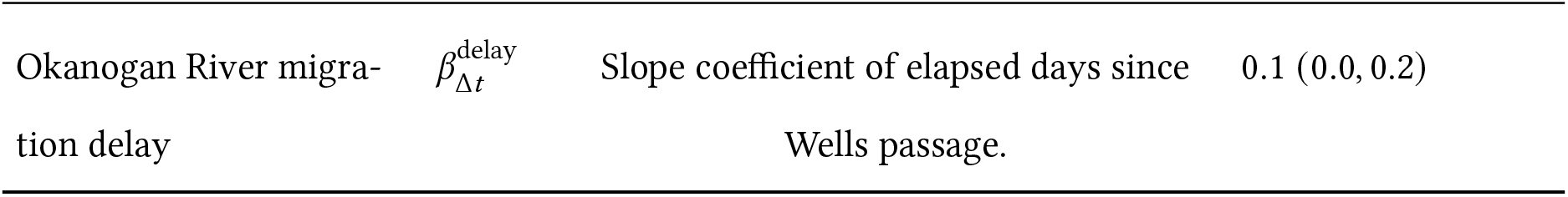
Coefficient estimates for logistic regressions describing Okanagan sockeye responses to environmental variables.

| Response variable | Parameter | Description | Est. (95% CI) |
|---|---|---|---|
| Lake Pateros holding | α<sup>hold</sup> | Population-level intercept | −3.7 (−4.0, −3.4) |
| Lake Pateros holding | β<sub>T̂</sub><sup>hold</sup> | Slope coefficient of rolling 4-day mean Okanogan River temperatures. | 1.2 (1.1, 1.2) |
| Lake Pateros holding | β<sub>F</sub><sup>hold</sup> | Slope coefficient of daily mean Okanogan River flows. | 1.1 (1.1, 1.2) |
| Lake Pateros holding | β<sub>Δt</sub><sup>hold</sup> | Slope coefficient of elapsed days since Wells passage. | 1.8 (1.7, 1.9) |
| Lake Pateros holding | β<sub>T̂Δt</sub><sup>hold</sup> | Interaction coefficient between *T̂* and *D*. | −0.26 (−0.28, −0.24) |
| Natural mortality | α<sup>mort</sup> | Population-level intercept | −10.1 (−10.7, −9.49) |
| Natural mortality | β<sub>T</sub><sup>mort</sup> | Slope coefficient of daily mean Okanogan River temperatures. | 1.3 (1.2, 1.4) |
| Natural mortality | β<sub>F</sub><sup>mort</sup> | Slope coefficient of daily mean Okanogan River flows. | −1.3 (−1.5, −1.1) |
| Natural mortality | β<sub>Δt</sub><sup>mort</sup> | Slope coefficient of elapsed days since Wells passage. | 0.1 (0.0, 0.2) |
| Okanogan River migra-tion delay | α<sup>delay</sup> | Population-level intercept | −0.6 (−1.0, −0.2) |
| Okanogan River migra-tion delay | β<sub>T</sub><sup>delay</sup> | Slope coefficient of daily mean Okanogan River temperatures. | 0.1 (0.0, 0.1) |
| Okanogan River migra-tion delay | β<sub>F</sub><sup>delay</sup> | Slope coefficient of daily mean Okanogan River flows. | 0.5 (0.4, 0.6) |
| Okanogan River migra-tion delay | β<sub>Δt</sub><sup>delay</sup> | Slope coefficient of elapsed days since Wells passage. | 0.1 (0.0, 0.2) |

Flow had more moderate effects on sockeye behaviour and mortality (Table 1). Okanogan River flows are highest in May and decline steadily during the sockeye migration season. When sockeye migration through Wells Dam peaks in early July, flows average 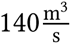 with a corresponding thalweg velocity of 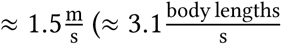; Figure S3; Nass et al. 2006). Flows typically reach their annual minima around 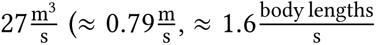; Figure S3 Nass et al. 2006) in early September. *P* (hold) increases from approximately 1% (1–2%) at 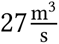 to approximately 9% (7–12%) at 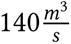; *P* (delay) also increases from approximately 31% (21–42%) to 49% (38–62%) between these two flows (Figure 3). Mortality conversely decreases as flows increase, from *P* (mort) ≈ 33% (26–73%) to ≈ 6% (3–9%) between 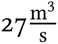 and 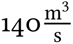 (based on a 24°C reference temperature; mortality is near 0% at the model intercept temperature of 19.9°C, regardless of flow; Figure 3).

### Forward simulation forecast quality

The median percentage forecast error for the forward simulations relative to observed annual total Zosel Dam sockeye passage for 2006–2010 and 2014–2016 is 20% (95% CI: 3–46%; Figure 4). Early in the season, *circa* 5 July, forecast uncertainty is high at approximately 150% (78–293%). However, by 25 July, expected uncertainty tapers to 30% (16–58%) and then declines gradually until stabilizing near 15% (8–30%) by 26 August (Figure 4B).

**Figure 4:**
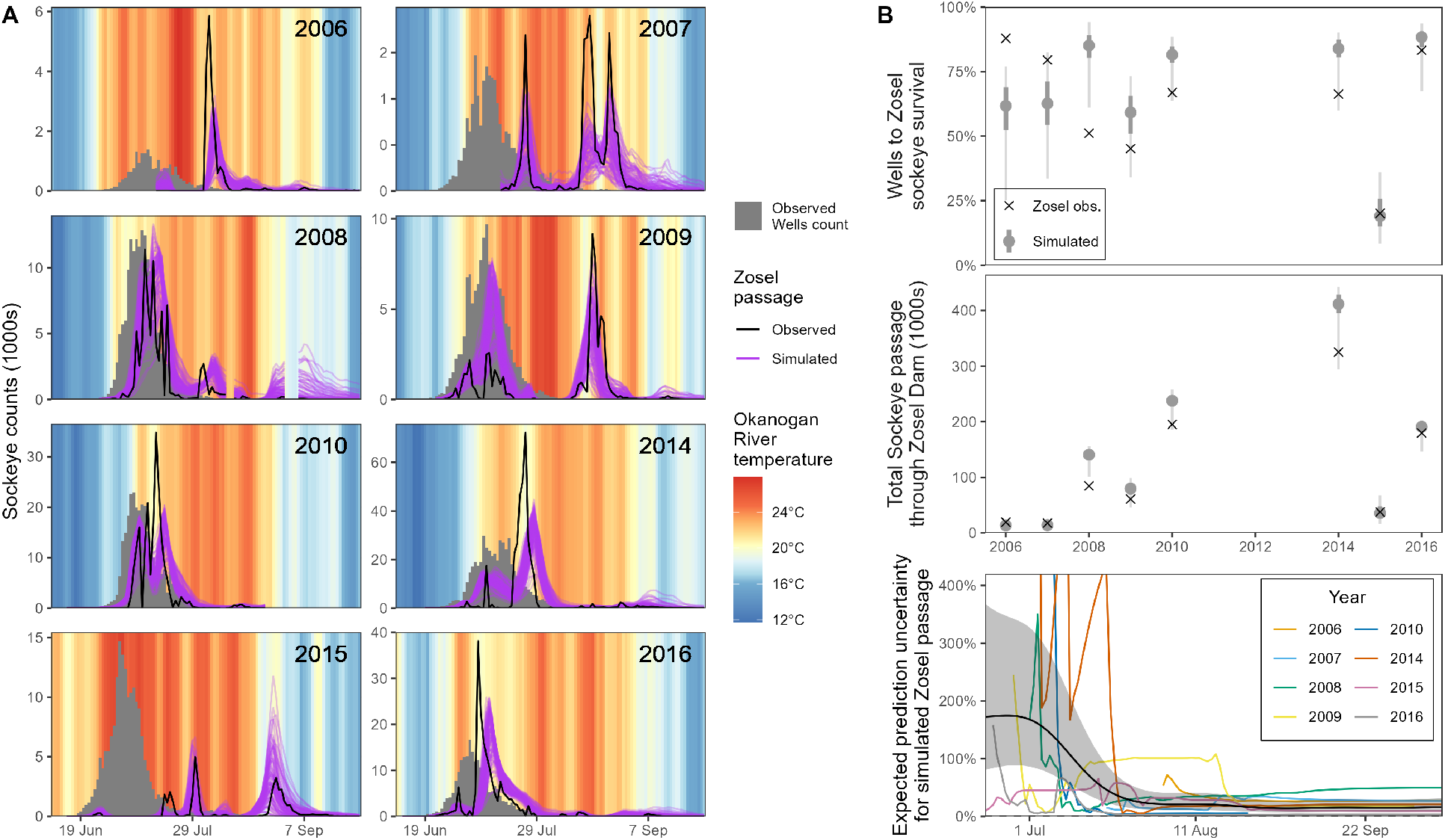
Simulated versus observed Zosel Dam sockeye passage for 2006–2010 and 2014–2016, years where observed data are most reliable. A) daily time series of observed Wells passage, Zosel passage, and Okanogan River temperatures overlaid with simulated Zosel passage data. Simulated data series are clipped to match gaps in the observed data. B) Annual total Wells to Zosel survival (top) and cumulative Zosel passage (middle) for all simulated years, with gray points, thick whiskers, and thin whiskers showing the median, interquartile range, and 95% CI of simulation outcomes, respectively. The bottom panel shows the GAMM median and 95% CI of daily expected forecast error, expressed relative to cumulative observed Zosel passage, with annual error time series shown as coloured lines. *Note*.—the y-axis scale in the bottom panel is capped at 400% but some daily values from 2010 and 2014 extend to 1800%.

### Simulation results

Results of population-level simulations for all years with available data are depicted in Figure 5. Comparing 2011 and 2015 illustrates the impact river temperatures can have on migration success in this system. In 2011, nearly all sockeye passing Wells Dam encountered Okanogan River temperatures below 20°C resulting in approximately 99% (96–99%) predicted survival to Zosel Dam. In 2015, Okanogan River temperatures exceeded 21°C from mid-June until the end of August, resulting in high vulnerability to harvest, high in-river moralities, and a predicted survival to Zosel Dam of only about 23% (19–25%; Figures 5 & 4B).

**Figure 5:**
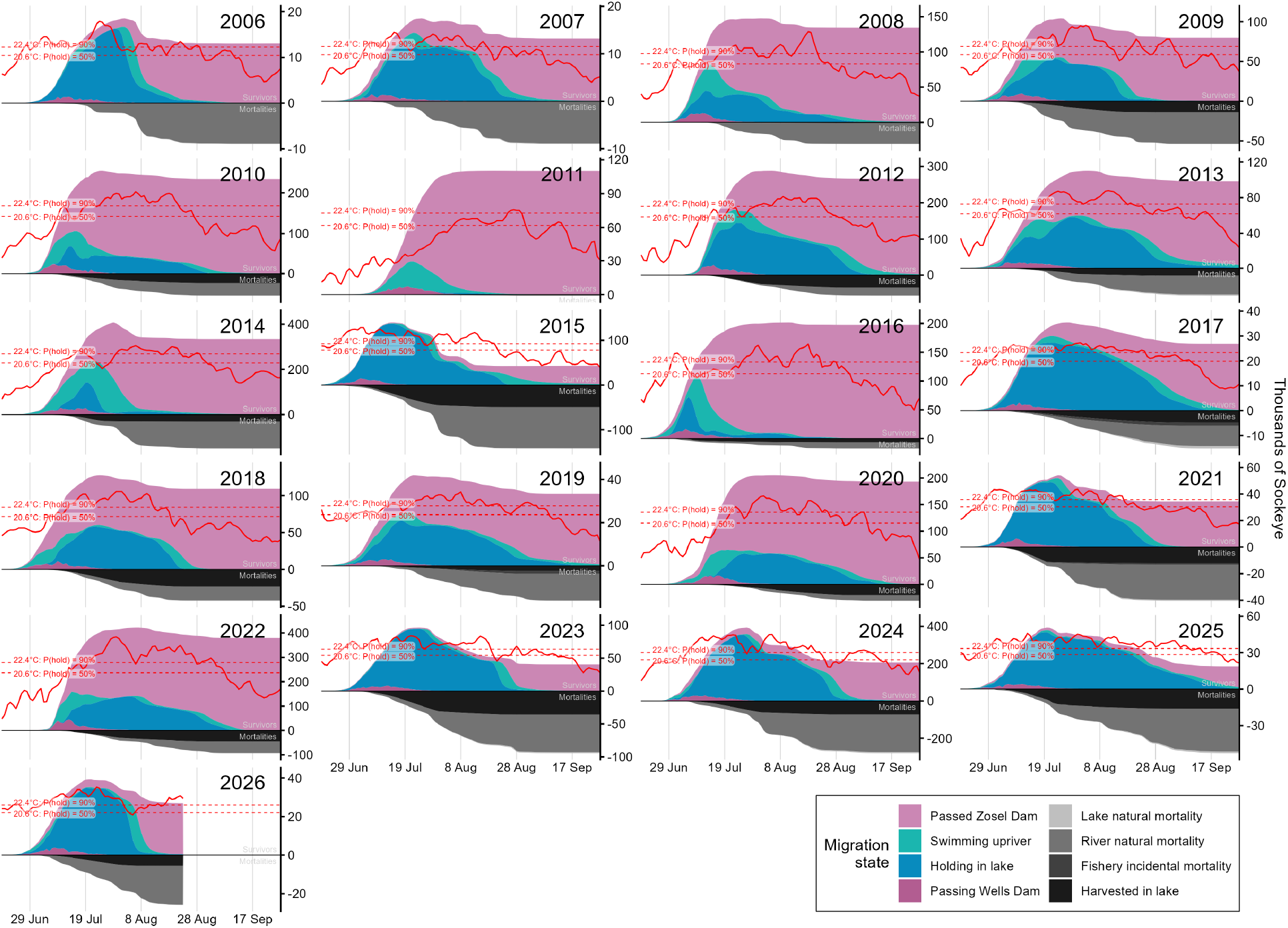
Simulated Okanagan sockeye migration between Wells and Zosel dams from 2006-2026, spanning all years with available environmental data. Daily values for each year and migration state are averaged over all posterior draws that were passed to the simulation. Red lines show the trajectory of daily Okanogan River temperatures and are plotted with respect to the red, dashed reference lines, not the right y-axis.

The simulations also reveal the possible perils associated with short-lived episodes of cooling river temperatures during peak migration. Most notably, in 2006, 2014, 2015, and 2024, river temperatures briefly dipped below *P* (hold) = 90% at 22.4°C while a large portion of the run was presumed to be holding in Lake Pateros, triggering significant influxes of migrants into the river who were quickly exposed to supra-optimal temperatures, apparently resulting in high-magnitude natural moralities (Figure 5).

## Discussion

We developed a forecasting framework that links Okanagan sockeye migration dynamics to daily river conditions. Leveraging a large dataset of PIT-tagged sockeye, we estimated functional relationships governing fish movement and mortality and found evidence that thermal tolerance declines with prolonged migration delays. When propagated through a population-level simulation, these relationships produced useful predictions of cumulative Zosel passage within one month of the first fish arriving at Wells Dam. These results highlight the strong effects of environmental conditions on salmon migration success, and showcase how combining individual movement and environmental data can enable key scientific advice for management decisions.

As temperatures warm, Okanogan sockeye cease migrating and mortality increases. The thermal thresholds we estimated for 50% daily probabilities of Okanagan sockeye holding and mortality—around 21°C and 25°C, respectively—align with previous estimates. A thermal barrier for sockeye entering the Okanogan River has been inferred at 21°C since at least the 1960s (Major and Mighell 1966) and 21°C was also identified as an inflection point in migration speeds for Columbia River chinook salmon (*O. tshawytscha*) (Goniea et al. 2006). Slightly lower thresholds *circa* 19°C have been reported for some Fraser River sockeye populations (Macdonald et al. 2010, Martins et al. 2011). These temperature thresholds likely reflect underlying physiological mechanisms. Returning adult salmon generally rely on aerobic metabolism to power upriver swimming, but also utilize anaerobic metabolism to overcome challenging rapids or other obstacles (Eliason et al. 2011). Water temperature affects salmon migration through its influence on aerobic scope (Mayer et al. 2024). Once ambient temperatures exceed a fish’s “pejus” threshold, aerobic capacity becomes increasingly limited (Pörtner and Knust 2007). Thermal barriers tend to be observed near salmon pejus temperatures, suggesting migration ceases when they can no longer swim upstream aerobically (Mayer et al. 2024).

Sockeye salmon populations are locally adapted to the environmental conditions encountered during migration (Crossin et al. 2004, Eliason et al. 2011). Variation in the response to flow and temperature between Okanogan and Fraser River sockeye may partly reflect differences in migration conditions. Our results showed that river flows have relatively modest effects on migration compared to water temperatures in the Okanogan River, while flow presents strong selection pressure on migration in the Fraser River (Macdonald et al. 2010, Eliason et al. 2011). The Columbia River and its tributaries, including the Okanogan, are extensively hydraulically regulated by dams and other flow mediation structures whereas the Fraser is relatively unregulated and presents more challenging swimming conditions for sockeye (Quinn et al. 1997, Hyatt et al. 2020, Wright et al. 2025). The flow data we extracted from 2005–2025 indicate Okanogan thalweg velocities rarely exceed 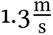 during sockeye migration while thalweg velocities average around 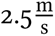 through the Fraser Canyon during sockeye migration (Wright et al. 2025). Similarly, the apparently higher holding threshold for Okanagan versus Fraser sockeye (≈ 21°C versus ≈ 19°C) likely reflects differences in migration thermal history. Okanagan sockeye are at the southern periphery of the species’ range and are already commonly facing stressful temperatures *>*21 °C, making them especially vulnerable to ongoing climate warming.

Thermal tolerances decreased with prolonged migration delays in our study. Specifically, the threshold temperature deterring entry into the Okanogan River decreased by approximately 1.5°C after fish spent one month holding in Lake Pateros. This pattern suggests that thermal tolerance declines over time, which may reflect a broader deterioration of physiological performance as migration and maturation progresses. Adult sockeye salmon cease feeding in the ocean, so upriver migration, sexual maturation and spawning behaviours are fueled entirely by finite endogenous energy stores. Previous work has reported that swimming performance declines throughout the migration with advancing sexual maturation in pink salmon (Williams and Brett 1987). Warm temperatures disproportionally reduced Fraser River sockeye salmon survival during the final states of migration, particularly among females (Martins et al. 2012). Nevertheless, the mechanisms underlying the apparent decline in thermal tolerance over time remain unclear and may include declining aerobic scope, energy depletion, reproductive maturation, and increased pathogen susceptibility (Whitney et al. 2016).

Both simulated and observed Zosel counts show that sockeye tend to push into the Okanogan River during brief cooling spells in July and August (when these occur) rather than holding until more reliable seasonal cooling kicks in around mid-September. This strategy may give them the best chances of reaching refuge in Osoyoos Lake before their energy reserves are exhausted, but could potentially predispose them to an ecological trap if temperatures quickly rebound while they are still ascending (Major and Mighell 1966). Under climate change, the thermal risks to Okanagan sockeye may therefore involve not only increases in average water temperatures, but also shifting duration and reliability of cooling windows. For example, wildfires are intensifying in North America (Curasi et al. 2024) and the resulting smoke can attenuate solar radiation, leading to temporary cooling in local streams (David et al. 2018). In 2026, the Okanogan River cooled more than 3°C between late July and early August, coincident with rapid growth of the nearby Sinlahekin Creek wildfire, providing early reprieve from the thermal barrier. Future fire activity may be similarly beneficial or harmful depending on the reliability of smoke coverage.

Given the complexities emerging from regional climate change, fishery managers require a sound framework for predicting the abundance and timing of Okanagan sockeye returning to Canada. The current assumption of a static 60% survival between Wells Dam and the Canadian border (Fisheries and Oceans Canada 2025) is insufficiently rigorous to meet the challenges arising from climate change. Our results show survival through the Okanogan River can range between ≈20–99%, depending on environmental conditions, and that an environmentally-conditioned mechanistic model can accurately predict survival and migration timing. However, the simulated Zosel counts are consistently smoother over time compared to the observed data, indicating that some sources of fine-scale variation remain unresolved. One plausible contributor is unmodeled social behaviour. Sockeye salmon are gregarious and tend to travel together in schools throughout their life cycle (Polyakov et al. 2022, Banks 1969, Berdahl et al. 2016). Berdahl et al. (2017) found that migration timing of sockeye ascending from Lake Aleknagik, AK, into Hansen Creek exhibited synchrony above and beyond what could be attributed to environmental triggers, and showed that a one-way decision copying mechanism could reproduce this synchrony mathematically. Uncertainty in model inputs, including harvest timing resolution, as well as other unmodeled processes may also contribute to discrepancies between predicted and observed counts. These limitations primarily affect the model’s ability to reproduce discrete daily arrivals rather than cumulative arrivals. Our forecast quality assessment showed that by late July, simulated Zosel arrivals constitute a defensible return-to-Canada prediction that could be used to inform adjusted harvest rates and opening decisions.

Okanagan sockeye are at a critical confluence between successful rehabilitation and mounting threats from climate change. Various fish passage, habitat quality, and flow management improvements, combined with a period of favourable marine conditions, allowed the population to rebound from its prior depressed state (Hyatt and Stockwell 2019, Alexander et al. 2024). Meanwhile, rising global temperatures are increasing the frequency and duration of thermal barrier events in the Okanogan River (Stiff and Thompson 2026), leading to elevated natural mortality rates and reduced reproductive potential of salmon that survive to reach the spawning grounds (Thompson et al. 2026). Our work provides detailed insights into the thermal tolerances of the population and a method to forecast the abundance returning to Canada. Such tools provide a basis for adaptive management strategies that can help mitigate the deleterious effects of climate change on Pacific salmon.

## Author contributions

P.T. and N.B. conceived the main ideas. N.B. led the data assembly, model development, analysis, and manuscript writing. P.T. and E.E. provided conceptual guidance and contributed to manuscript writing and editing.

## Acknowledgments

J. Fryer (Columbia River Inter-Tribal Fish Commission) and E. McGrath (Okanagan Nation Alliance) provided helpful background information on Columbia and Okanagan PIT tagging programs. C. Jackson (Washington Department of Fish and Wildlife) provided monthly Lake Pateros recreational kept and released catch data. K. Ryde (Fisheries and Oceans Canada) provided annual Lake Pateros total kept catch data. We used Microsoft’s Copilot Artificial Intelligence (AI) model (GPT-5, 2026) to draft some of the supporting R and Stan code. We thoroughly reviewed and revised all AI-generated content before incorporating it.

## Data availability

Supporting data and code are available from [Zenodo repository TBC].

## Supporting Information

### Okanogan River and Lake Pateros water temperature time series imputation

We used water temperatures recorded at U.S. Geological Survey (USGS) monitoring stations near Malott and Wells Dam, WA, as indices for water temperatures in the Okanogan River and Lake Pateros, respectively (USGS station IDs: 12447200 and 12450650, respectively). However, both stations have periods with missing data, including during the sockeye migration season, requiring data imputation to produce the complete daily time series needed for the Hidden Markov process model. Fortunately, monitoring stations with more complete records are situated upstream from both locations, which provided a strong basis for informative imputation in both time series.

We used 1-day-lagged water temperatures recorded by the U.S. Army Corps of Engineers monitoring station at the Chief Joseph Dam tailrace (at the head of Lake Pateros) as a predictor to impute Wells Dam (Lake Pateros) water temperatures 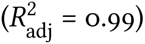. Exploratory plots revealed sporadic daily temperature values that deviated extremely from neighboring values and from the seasonal thermograph in both time series, so we removed these observations before proceeding with imputation (18 Wells Dam daily observations and 11 Chief Joseph Dam daily observations). We also included same-day water temperature observations from the Malott time series in the Lake Pateros imputation model to account for the influence of the Okanogan River, which confluences into Lake Pateros downstream from Chief Joseph Dam. The full imputation model was a generalized additive model (GAM) fitting Wells Dam temperatures as a function of a smooth seasonal trend (day of year), standardized 1-day-lagged Chief Joseph Dam temperatures, and standardized same-day Malott temperatures, with additional smooth-by-predictor terms permitting the influence of each temperature series to vary nonlinearly through time. Model smoothness was estimated using restricted maximum likelihood (REML), and shrinkage penalties were applied to minimize overfitting. When one or both upstream temperature series were unavailable, reduced models containing the available predictor or seasonal trend alone were used for imputation (Figure S1).

We used same-day water temperatures recorded at the USGS monitoring stations near Oroville on the Okanogan River and on the Similkameen River (USGS station IDs: 12439500 and 12442500, respectively) to impute missing Malott water temperatures. The final imputation model was a linear mixed-effects model fitting daily minimum, mean, and maximum Malott temperatures as functions of standardized same-day temperatures at Oroville and Similkameen and a quadratic seasonal trend based on day of the year, with separate fixed-effect coefficients estimated for each temperature statistic (minimum, mean, and maximum). The model also included a random intercept for year and independent year-specific random slopes for the Oroville and Similkameen predictors, allowing both baseline temperatures and relationships among stations to vary among years. The final model was selected based on leave-one-year-out cross-validation and fitted using REML. When one or both upstream temperature series were unavailable, reduced models containing the available predictor or seasonal trend alone were used for imputation (Figure S2).

**Figure S1:**
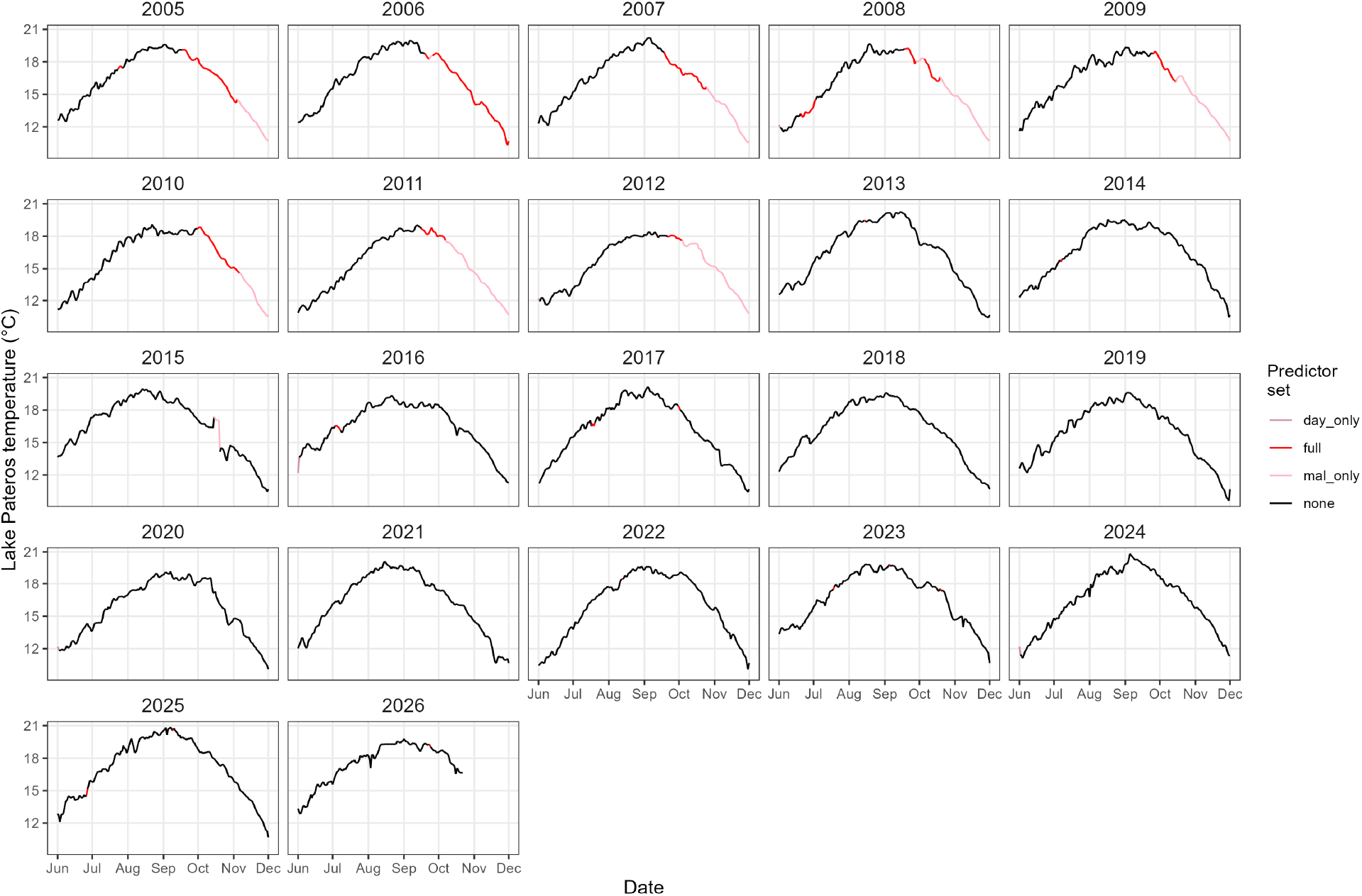
Lake Pateros water temperature time series from USGS monitoring station 12450650 at Wells Dam, WA. Black lines show observed daily mean temperatures. Red and pink lines show imputed daily temperatures according to which predictor set was available and used to inform the imputed values. Values used as predictors in the interpolation model included day of year, 1-day-lagged water temperatures recorded by the U.S. Army Corps of Engineers monitoring station at the Chief Joseph Dam tailrace, and same-day water temperatures recorded at USGS monitoring station 12447200 at Malott, WA. Interpolated values were informed by as many of these predictors as possible on each day.

**Figure S2:**
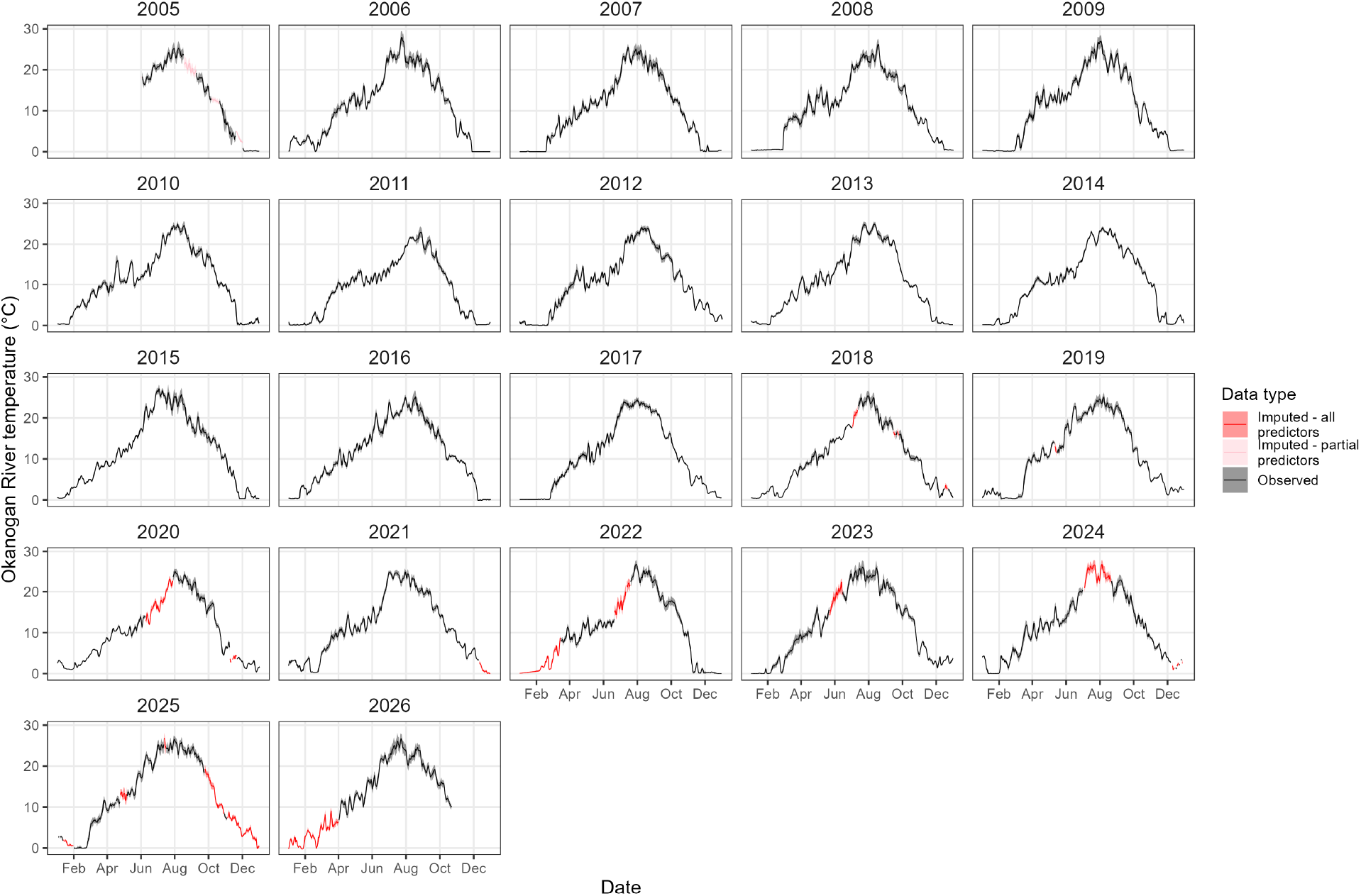
Okanogan River water temperature time series from USGS monitoring station 12447200 at Malott, WA. Black lines and grey shaded areas show observed daily mean and minimum–maximum temperatures, respectively. Red and pink lines and shaded areas show imputed daily mean and minimum–maximum temperatures, respectively. Values used as predictors in the interpolation model included day of year and same-day temperatures recorded upstream at USGS monitoring stations 12439500 and 12442500 on the upper Okanogan and Similkameen rivers, respectively. Interpolated values were informed by as many of these predictors as possible on each day.

**Figure S3:**
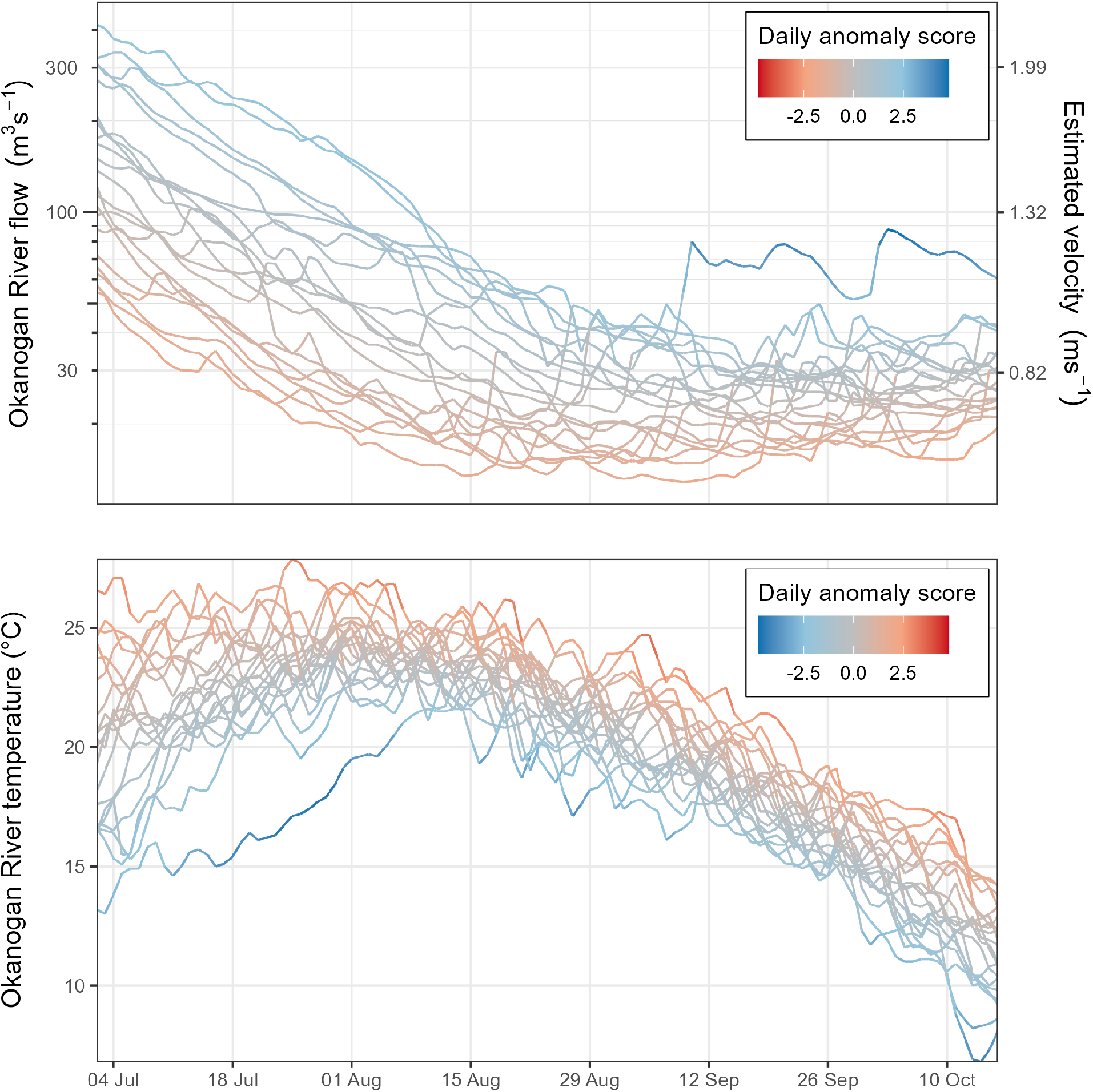
Okanagan River temperature and flow time series during the sockeye migration seasons of 2005–2025. Each line shows the time series for one year. Lines are coloured according to daily anomaly (z) score values. Note that the colour palette is reversed between the two panels, conveying the general idea that cooler temperatures and higher flows constitute better conditions for sockeye in this watershed. The right y-axis in the top panel shows the approximate thalweg velocities based on flow; these values were estimated using monotonic functions reported in Nass et al. (2006).

### HMM latent state definitions

All fish were initialized alive at Wells Dam and then propagated into the following latent states:

**Table S1:** Latent states in the Wells-to-Zosel migration model.

| State | Description |
|---|---|
| 1 | Holding in Lake Pateros |
| 2 | Direct pre-OKL transit without prior holding |
| 3 | Pre-OKL transit after Lake Pateros holding |
| 4 | Post-OKL migration delay |
| 5 | Final migration day toward Zosel |
| 6 | Natural mortality or fishing-related incidental mortality |
| 7 | Reported kept harvest |
| 8 | Arrived at Zosel |

### Prior Values

Priors used in the HMM + harvest recursion.

**Table S2:**
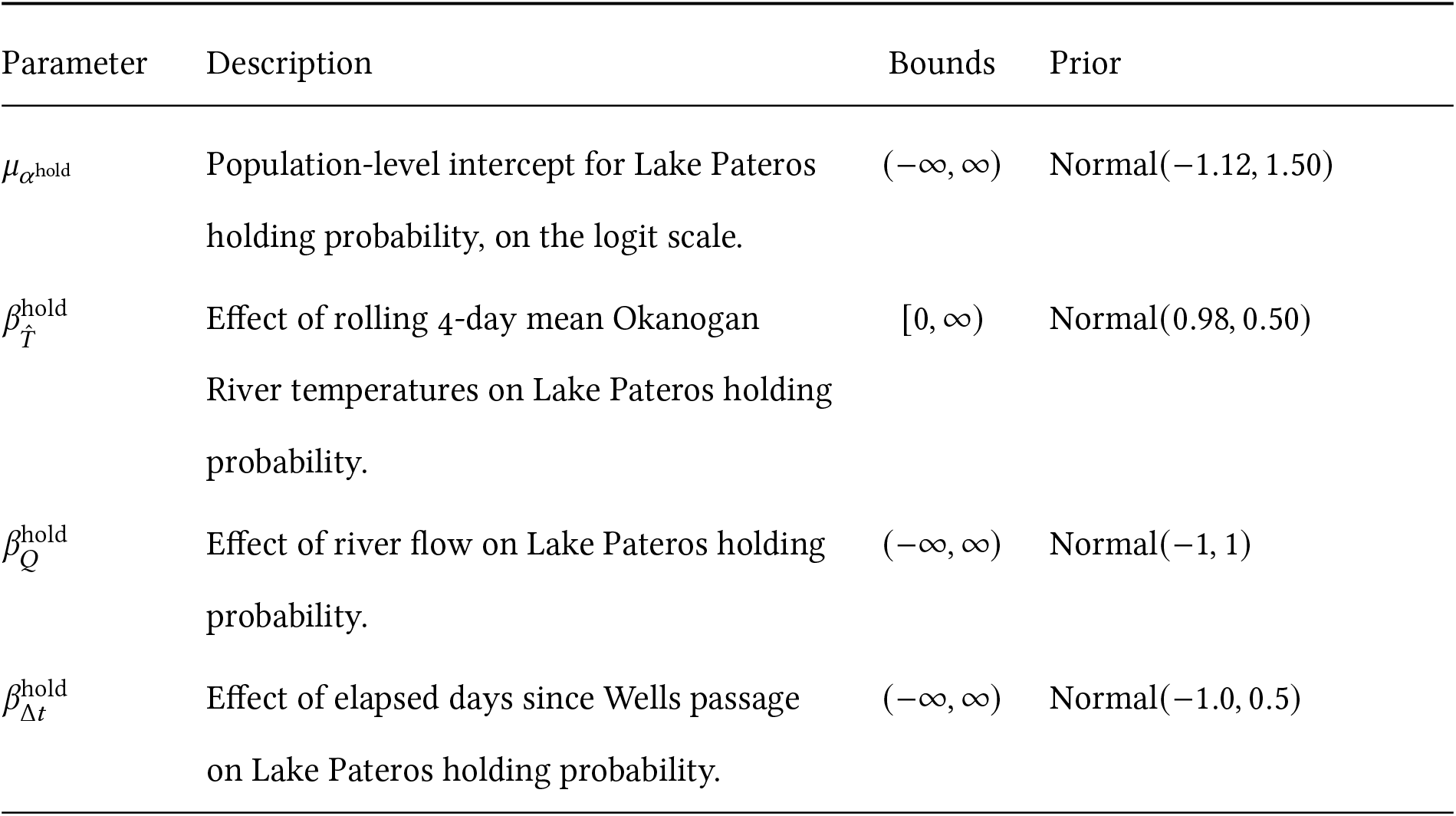
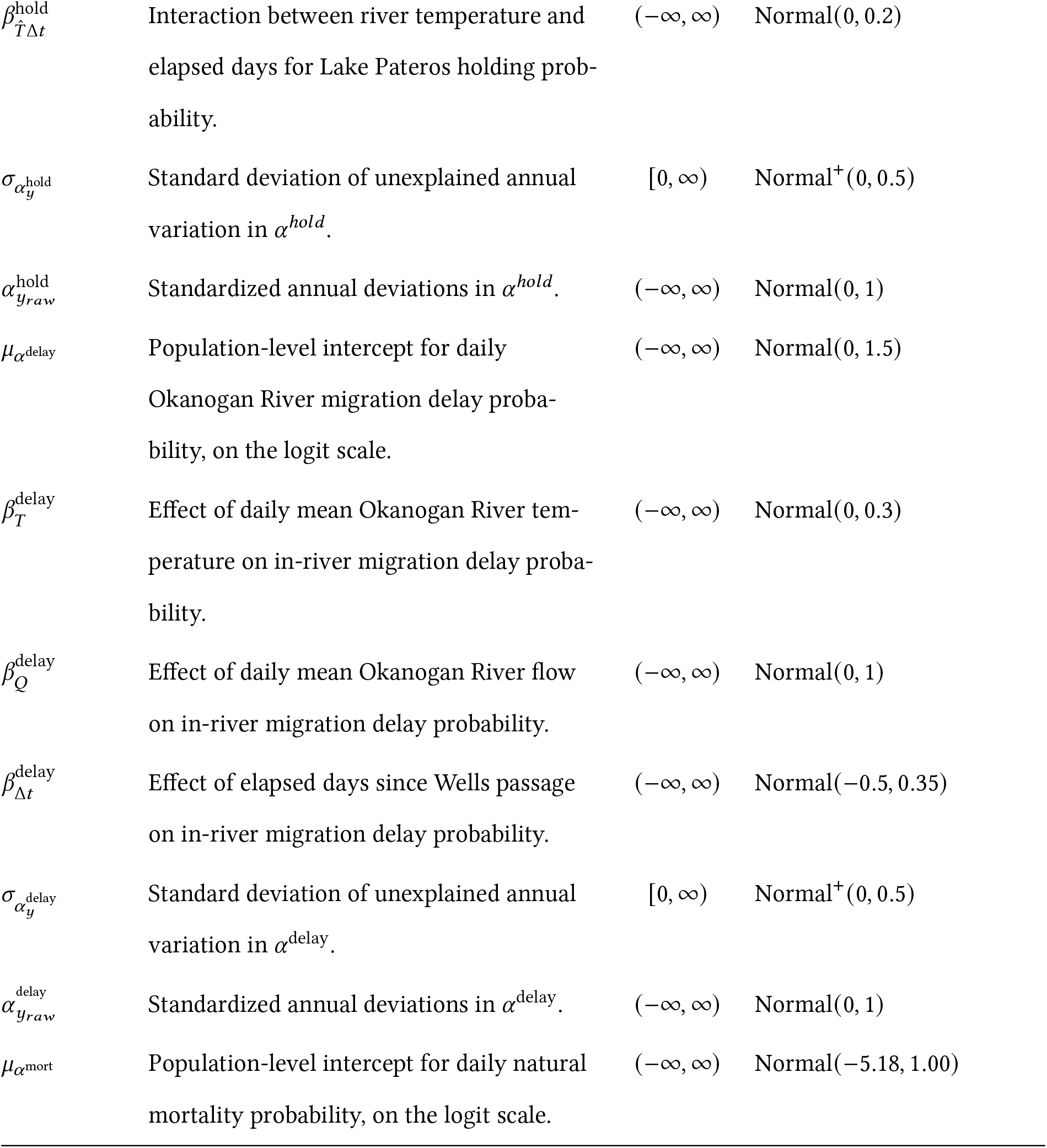
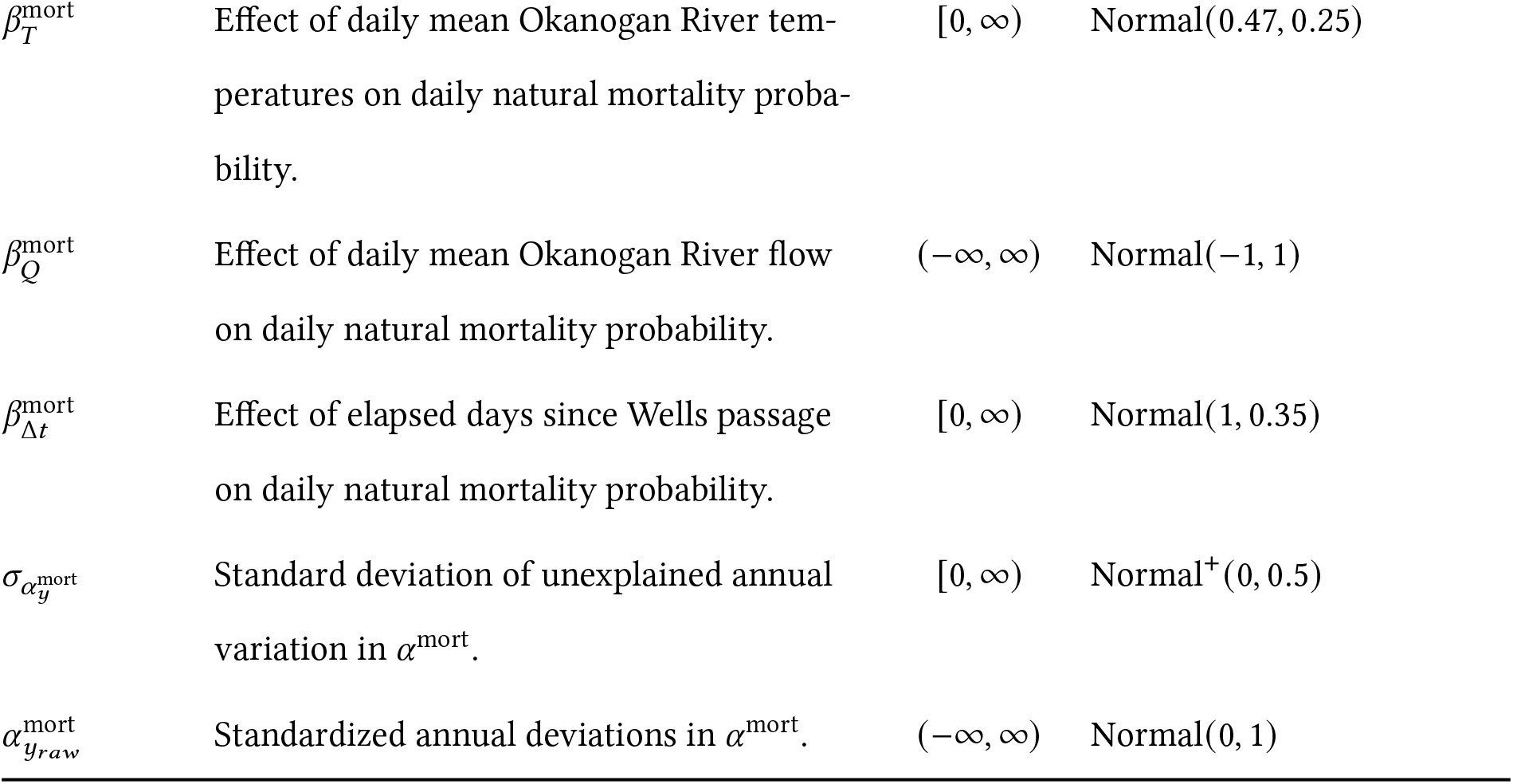
Prior distributions for logistic regressions describing Sockeye responses to environmental variables.

| Parameter | Description | Bounds | Prior |
|---|---|---|---|
| μ<sub>α<sup>hold</sup></sub> | Population-level intercept for Lake Pateros holding probability, on the logit scale. | (−∞, ∞) | Normal(−1.12, 1.50) |
| β<sup>hold</sup><sub>T̂</sub> | Effect of rolling 4-day mean Okanogan River temperatures on Lake Pateros holding probability. | [0, ∞) | Normal(0.98, 0.50) |
| β<sup>hold</sup><sub>Q</sub> | Effect of river flow on Lake Pateros holding probability. | (−∞, ∞) | Normal(−1, 1) |
| β<sup>hold</sup><sub>Δt</sub> | Effect of elapsed days since Wells passage on Lake Pateros holding probability. | (−∞, ∞) | Normal(−1.0, 0.5) |
| β<sup>hold</sup><sub>T̂Δt</sub> | Interaction between river temperature and elapsed days for Lake Pateros holding prob-ability. | (−∞, ∞) | Normal(0, 0.2) |
| σ<sub>α<sub>y</sub><sup>hold</sup></sub> | Standard deviation of unexplained annual variation in α<sup>*hold*</sup>. | [0, ∞) | Normal<sup>+</sup>(0, 0.5) |
| α<sup>hold</sup><sub>y<sub>raw</sub></sub> | Standardized annual deviations in α<sup>*hold*</sup>. | (−∞, ∞) | Normal(0, 1) |
| μ<sub>α<sup>delay</sup></sub> | Population-level intercept for daily Okanogan River migration delay proba-bility, on the logit scale. | (−∞, ∞) | Normal(0, 1.5) |
| β<sup>delay</sup><sub>T</sub> | Effect of daily mean Okanogan River tem-perature on in-river migration delay proba-bility. | (−∞, ∞) | Normal(0, 0.3) |
| β<sup>delay</sup><sub>Q</sub> | Effect of daily mean Okanogan River flow on in-river migration delay probability. | (−∞, ∞) | Normal(0, 1) |
| β<sup>delay</sup><sub>Δt</sub> | Effect of elapsed days since Wells passage on in-river migration delay probability. | (−∞, ∞) | Normal(−0.5, 0.35) |
| σ<sub>α<sub>y</sub><sup>delay</sup></sub> | Standard deviation of unexplained annual variation in α<sup>delay</sup>. | [0, ∞) | Normal<sup>+</sup>(0, 0.5) |
| α<sup>delay</sup><sub>y<sub>raw</sub></sub> | Standardized annual deviations in α<sup>delay</sup>. | (−∞, ∞) | Normal(0, 1) |
| μ<sub>α<sup>mort</sup></sub> | Population-level intercept for daily natural mortality probability, on the logit scale. | (−∞, ∞) | Normal(−5.18, 1.00) |
| β<sup>mort</sup><sub>T</sub> | Effect of daily mean Okanogan River tem-peratures on daily natural mortality proba-bility. | [0, ∞) | Normal(0.47, 0.25) |
| β<sup>mort</sup><sub>Q</sub> | Effect of daily mean Okanogan River flow on daily natural mortality probability. | (−∞, ∞) | Normal(−1, 1) |
| β<sup>mort</sup><sub>Δt</sub> | Effect of elapsed days since Wells passage on daily natural mortality probability. | [0, ∞) | Normal(1, 0.35) |
| σ<sub>α<sub>y</sub><sup>mort</sup></sub> | Standard deviation of unexplained annual variation in α<sup>mort</sup>. | [0, ∞) | Normal<sup>+</sup>(0, 0.5) |
| α<sup>mort</sup><sub>y<sub>raw</sub></sub> | Standardized annual deviations in α<sup>mort</sup>. | (−∞, ∞) | Normal(0, 1) |

**Table S3:**
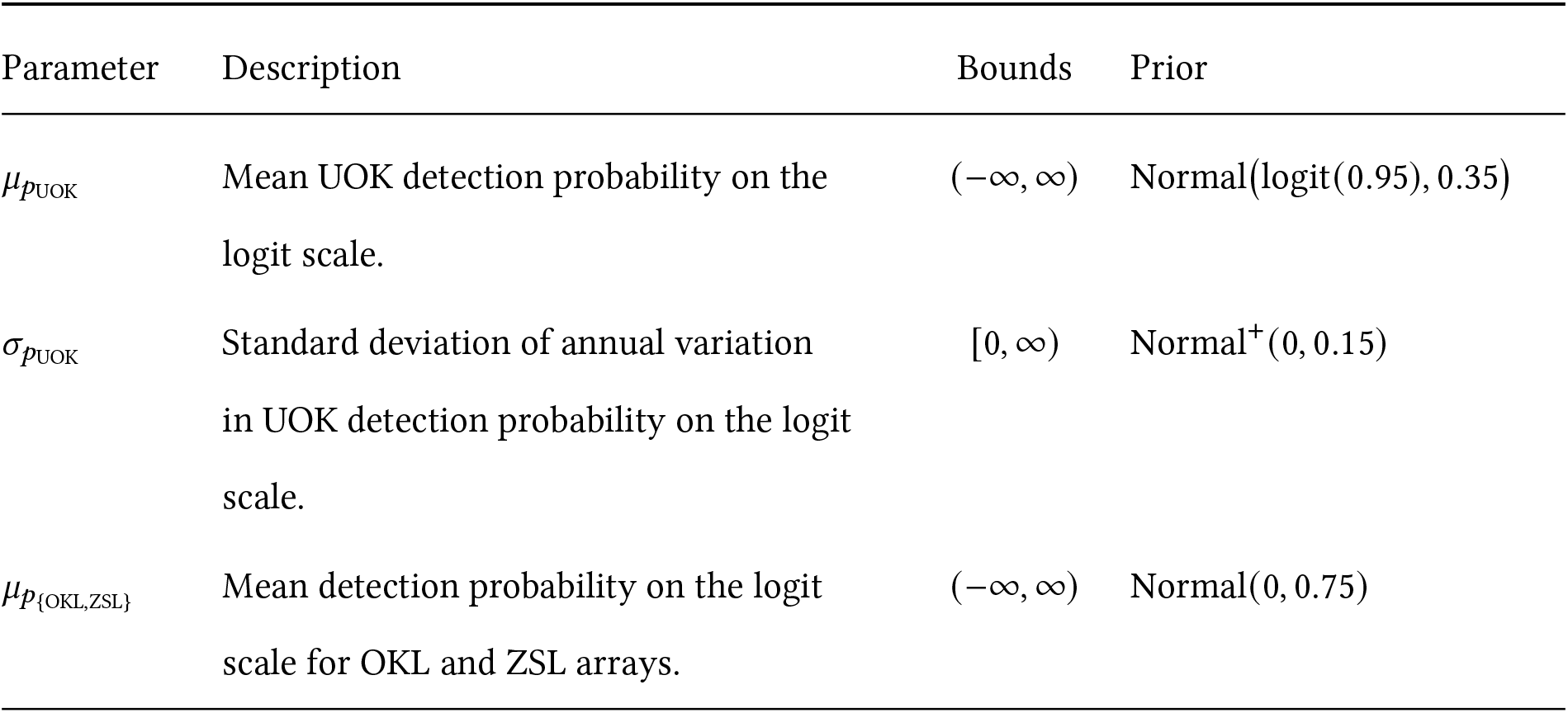
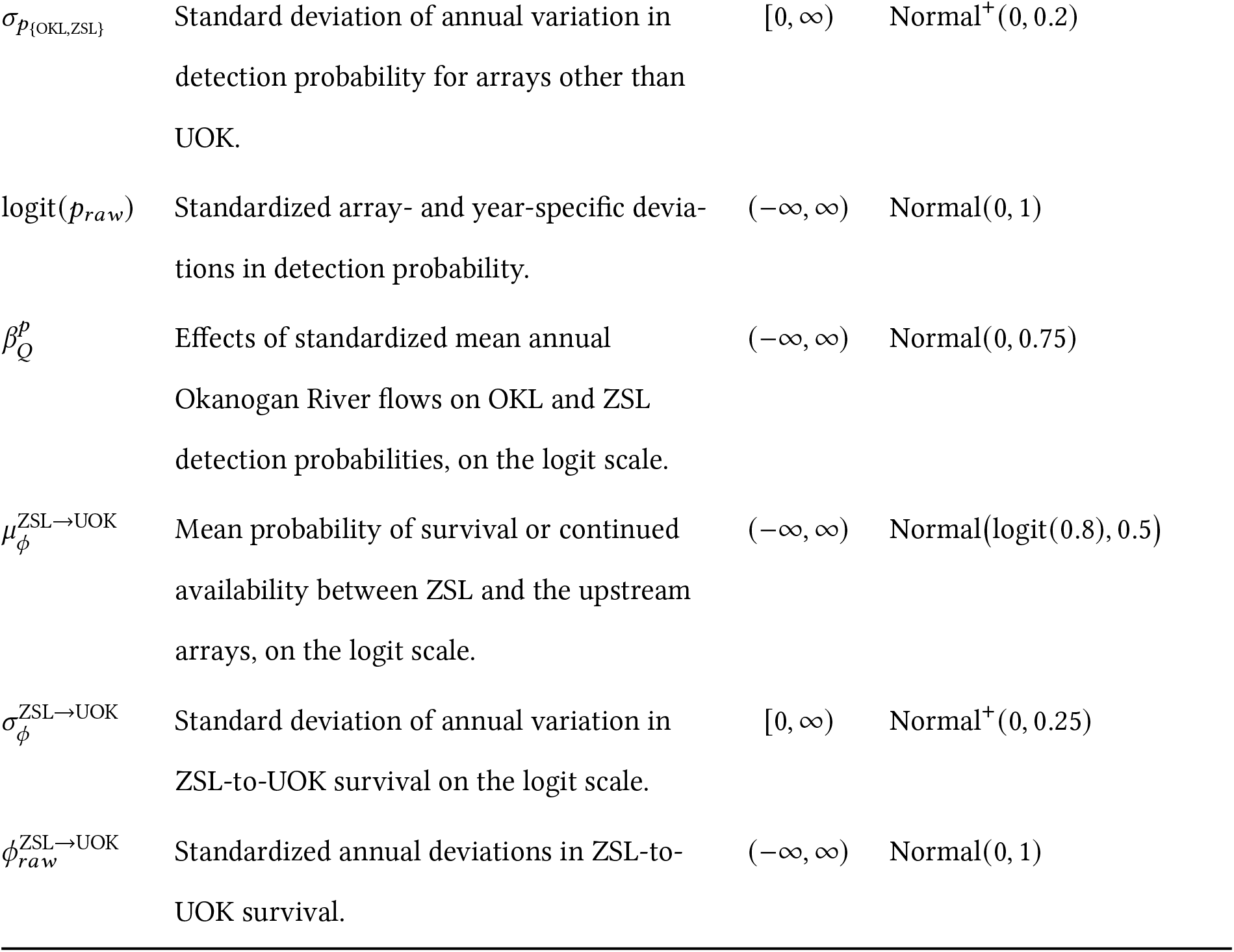
Prior distributions for Cormack-Jolly-Seber PIT tag observation sub-model.

| Parameter | Description | Bounds | Prior |
|---|---|---|---|
| μ<sub>p<sub>UOK</sub></sub> | Mean UOK detection probability on the logit scale. | (−∞, ∞) | Normal(logit(0.95), 0.35) |
| σ<sub>p<sub>UOK</sub></sub> | Standard deviation of annual variation in UOK detection probability on the logit scale. | [0, ∞) | Normal<sup>+</sup>(0, 0.15) |
| μ<sub>p<sub>{OKL,ZSL}</sub></sub> | Mean detection probability on the logit scale for OKL and ZSL arrays. | (−∞, ∞) | Normal(0, 0.75) |
| σ<sub>p<sub>{OKL,ZSL}</sub></sub> | Standard deviation of annual variation in detection probability for arrays other than UOK. | [0, ∞) | Normal<sup>+</sup>(0, 0.2) |
| logit(p<sub>raw</sub>) | Standardized array- and year-specific devia-tions in detection probability. | (−∞, ∞) | Normal(0, 1) |
| β<sup>p</sup><sub>Q</sub> | Effects of standardized mean annual Okanogan River flows on OKL and ZSL detection probabilities, on the logit scale. | (−∞, ∞) | Normal(0, 0.75) |
| μ<sup>ZSL→UOK</sup><sub>ϕ</sub> | Mean probability of survival or continued availability between ZSL and the upstream arrays, on the logit scale. | (−∞, ∞) | Normal(logit(0.8), 0.5) |
| σ<sup>ZSL→UOK</sup><sub>ϕ</sub> | Standard deviation of annual variation in ZSL-to-UOK survival on the logit scale. | [0, ∞) | Normal<sup>+</sup>(0, 0.25) |
| ϕ<sup>ZSL→UOK</sup><sub>raw</sub> | Standardized annual deviations in ZSL-to-UOK survival. | (−∞, ∞) | Normal(0, 1) |

**Table S4:**
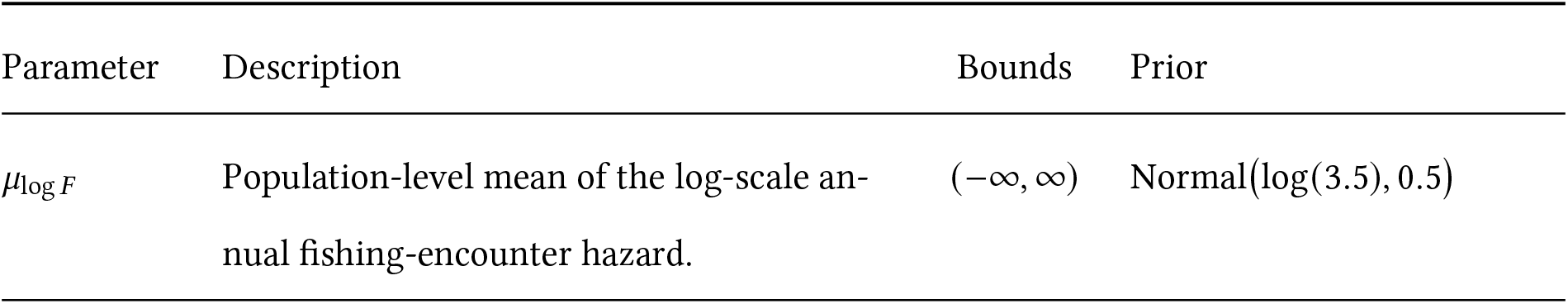
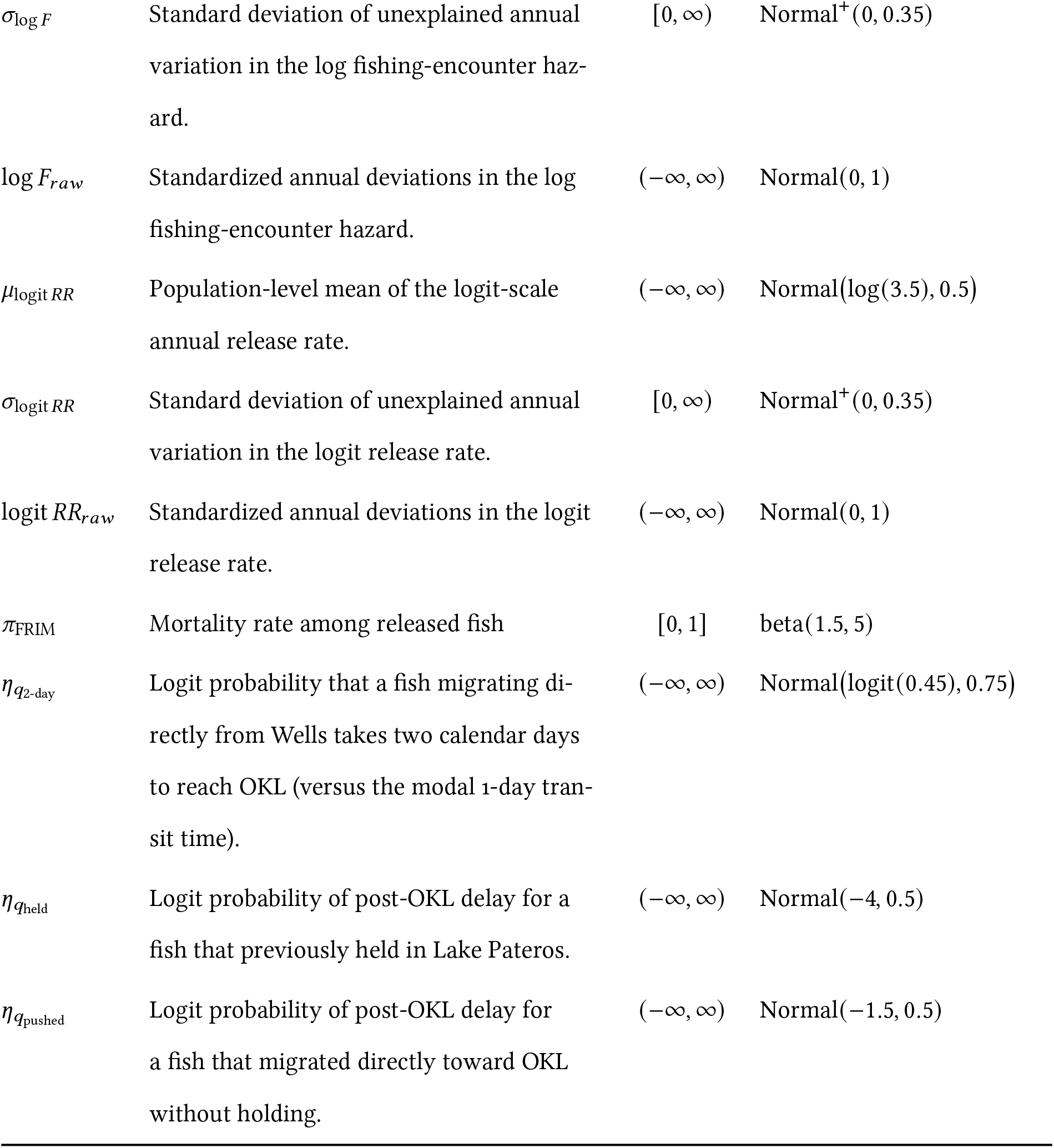
Prior distributions for additional parameters in the individual-(PIT) and population-level likelihoods.

| Parameter | Description | Bounds | Prior |
|---|---|---|---|
| μ<sub>log F</sub> | Population-level mean of the log-scale an-nual fishing-encounter hazard. | (−∞, ∞) | Normal(log(3.5), 0.5) |
| σ<sub>log F</sub> | Standard deviation of unexplained annual variation in the log fishing-encounter haz-ard. | [0, ∞) | Normal<sup>+</sup>(0, 0.35) |
| log F<sub>raw</sub> | Standardized annual deviations in the log fishing-encounter hazard. | (−∞, ∞) | Normal(0, 1) |
| μ<sub>logit RR</sub> | Population-level mean of the logit-scale annual release rate. | (−∞, ∞) | Normal(log(3.5), 0.5) |
| σ<sub>logit RR</sub> | Standard deviation of unexplained annual variation in the logit release rate. | [0, ∞) | Normal<sup>+</sup>(0, 0.35) |
| logit RR<sub>raw</sub> | Standardized annual deviations in the logit release rate. | (−∞, ∞) | Normal(0, 1) |
| π<sub>FRIM</sub> | Mortality rate among released fish | [0, 1] | beta(1.5, 5) |
| η<sub>q<sub>2-day</sub></sub> | Logit probability that a fish migrating di-rectly from Wells takes two calendar days to reach OKL (versus the modal 1-day tran-sit time). | (−∞, ∞) | Normal(logit(0.45), 0.75) |
| η<sub>q<sub>held</sub></sub> | Logit probability of post-OKL delay for a fish that previously held in Lake Pateros. | (−∞, ∞) | Normal(−4, 0.5) |
| η<sub>q<sub>pushed</sub></sub> | Logit probability of post-OKL delay for a fish that migrated directly toward OKL without holding. | (−∞, ∞) | Normal(−1.5, 0.5) |

### HMM transition logic and equations

**Table S5:**
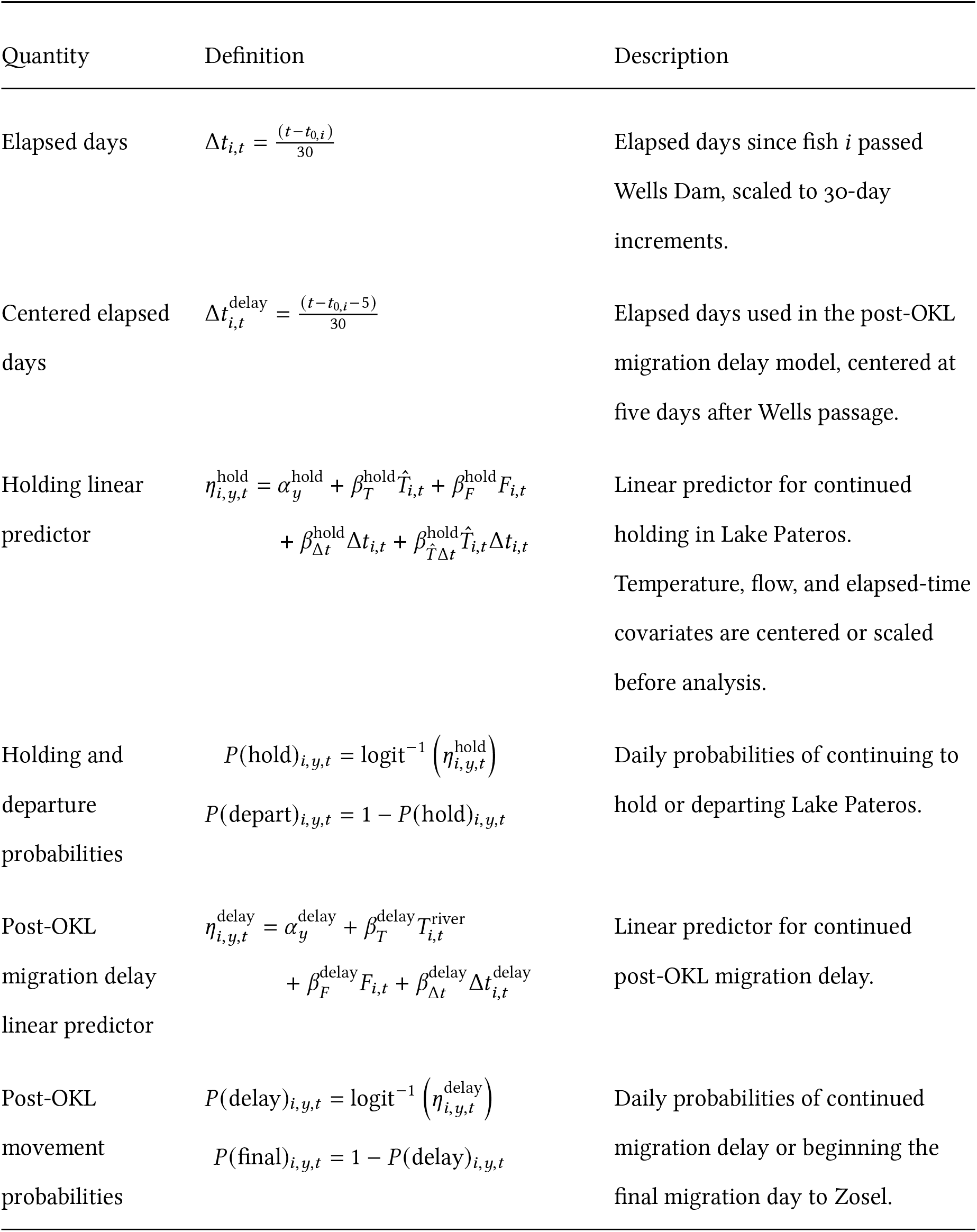
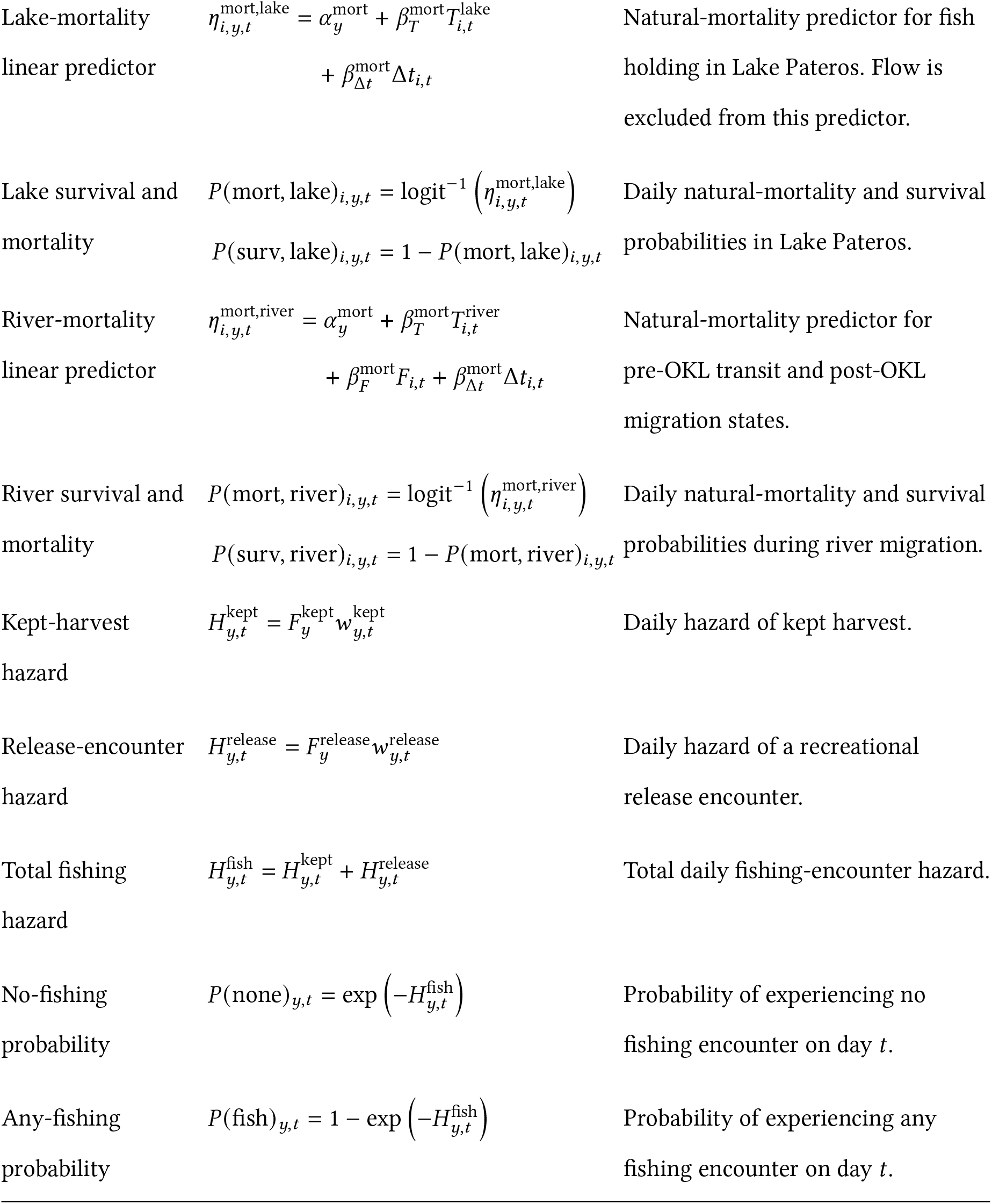
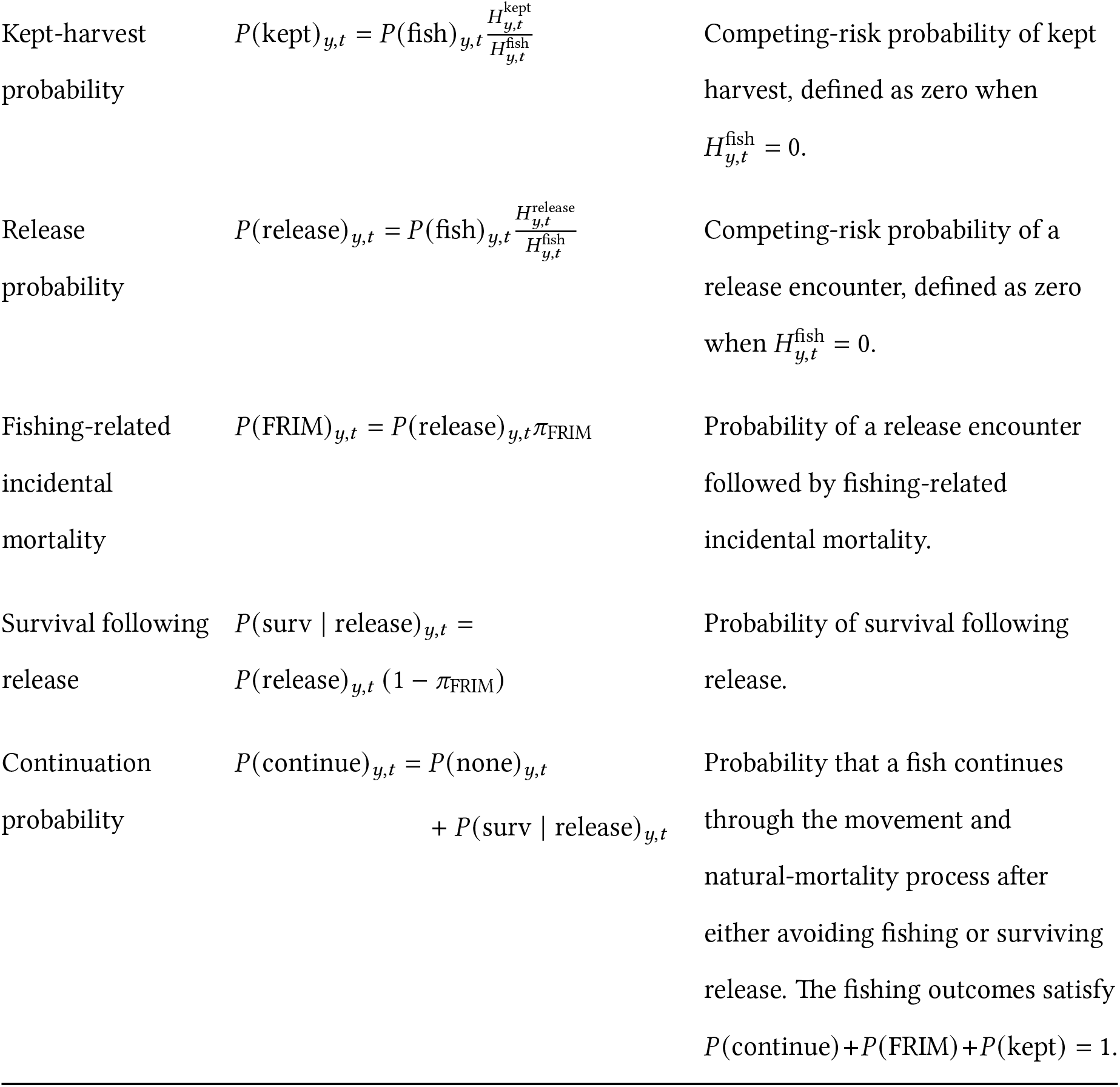
Notation and shared probability functions used in the state-transition model.

| Quantity | Definition | Description |
|---|---|---|
| Elapsed days | $\Delta t_{i,t} = \frac{(t - t_{0,i})}{30}$ | Elapsed days since fish *i* passed Wells Dam, scaled to 30-day increments. |
| Centered elapsed days | $\Delta t_{i,t}^{\text{delay}} = \frac{(t - t_{0,i} - 5)}{30}$ | Elapsed days used in the post-OKL migration delay model, centered at five days after Wells passage. |
| Holding linear predictor | $\eta_{i,y,t}^{\text{hold}} = \alpha_y^{\text{hold}} + \beta_T^{\text{hold}} \hat{T}_{i,t} + \beta_F^{\text{hold}} F_{i,t}$<br>$+ \beta_{\Delta t}^{\text{hold}} \Delta t_{i,t} + \beta_{\hat{T}\Delta t}^{\text{hold}} \hat{T}_{i,t} \Delta t_{i,t}$ | Linear predictor for continued holding in Lake Pateros. Temperature, flow, and elapsed-time covariates are centered or scaled before analysis. |
| Holding and departure probabilities | $P(\text{hold})_{i,y,t} = \text{logit}^{-1}\left(\eta_{i,y,t}^{\text{hold}}\right)$<br>$P(\text{depart})_{i,y,t} = 1 - P(\text{hold})_{i,y,t}$ | Daily probabilities of continuing to hold or departing Lake Pateros. |
| Post-OKL migration delay linear predictor | $\eta_{i,y,t}^{\text{delay}} = \alpha_y^{\text{delay}} + \beta_T^{\text{delay}} T_{i,t}^{\text{river}}$<br>$+ \beta_F^{\text{delay}} F_{i,t} + \beta_{\Delta t}^{\text{delay}} \Delta t_{i,t}^{\text{delay}}$ | Linear predictor for continued post-OKL migration delay. |
| Post-OKL movement probabilities | $P(\text{delay})_{i,y,t} = \text{logit}^{-1}\left(\eta_{i,y,t}^{\text{delay}}\right)$<br>$P(\text{final})_{i,y,t} = 1 - P(\text{delay})_{i,y,t}$ | Daily probabilities of continued migration delay or beginning the final migration day to Zosel. |
| Lake-mortality linear predictor | $\eta_{i,y,t}^{\text{mort,lake}} = \alpha_y^{\text{mort}} + \beta_T^{\text{mort}} T_{i,t}^{\text{lake}}$<br>$+ \beta_{\Delta t}^{\text{mort}} \Delta t_{i,t}$ | Natural-mortality predictor for fish holding in Lake Pateros. Flow is excluded from this predictor. |
| Lake survival and mortality | $P(\text{mort, lake})_{i,y,t} = \text{logit}^{-1}\left(\eta_{i,y,t}^{\text{mort,lake}}\right)$<br>$P(\text{surv, lake})_{i,y,t} = 1 - P(\text{mort, lake})_{i,y,t}$ | Daily natural-mortality and survival probabilities in Lake Pateros. |
| River-mortality linear predictor | $\eta_{i,y,t}^{\text{mort,river}} = \alpha_y^{\text{mort}} + \beta_T^{\text{mort}} T_{i,t}^{\text{river}}$<br>$+ \beta_F^{\text{mort}} F_{i,t} + \beta_{\Delta t}^{\text{mort}} \Delta t_{i,t}$ | Natural-mortality predictor for pre-OKL transit and post-OKL migration states. |
| River survival and mortality | $P(\text{mort, river})_{i,y,t} = \text{logit}^{-1}\left(\eta_{i,y,t}^{\text{mort,river}}\right)$<br>$P(\text{surv, river})_{i,y,t} = 1 - P(\text{mort, river})_{i,y,t}$ | Daily natural-mortality and survival probabilities during river migration. |
| Kept-harvest hazard | $H_{y,t}^{\text{kept}} = F_y^{\text{kept}} w_{y,t}^{\text{kept}}$ | Daily hazard of kept harvest. |
| Release-encounter hazard | $H_{y,t}^{\text{release}} = F_y^{\text{release}} w_{y,t}^{\text{release}}$ | Daily hazard of a recreational release encounter. |
| Total fishing hazard | $H_{y,t}^{\text{fish}} = H_{y,t}^{\text{kept}} + H_{y,t}^{\text{release}}$ | Total daily fishing-encounter hazard. |
| No-fishing probability | $P(\text{none})_{y,t} = \exp\left(-H_{y,t}^{\text{fish}}\right)$ | Probability of experiencing no fishing encounter on day *t*. |
| Any-fishing probability | $P(\text{fish})_{y,t} = 1 - \exp\left(-H_{y,t}^{\text{fish}}\right)$ | Probability of experiencing any fishing encounter on day *t*. |
| Kept-harvest probability | $P(\text{kept})_{y,t} = P(\text{fish})_{y,t} \frac{H_{y,t}^{\text{kept}}}{H_{y,t}^{\text{fish}}}$ | Competing-risk probability of kept harvest, defined as zero when $H_{y,t}^{\text{fish}} = 0$. |
| Release probability | $P(\text{release})_{y,t} = P(\text{fish})_{y,t} \frac{H_{y,t}^{\text{release}}}{H_{y,t}^{\text{fish}}}$ | Competing-risk probability of a release encounter, defined as zero when $H_{y,t}^{\text{fish}} = 0$. |
| Fishing-related incidental mortality | $P(\text{FRIM})_{y,t} = P(\text{release})_{y,t} \pi_{\text{FRIM}}$ | Probability of a release encounter followed by fishing-related incidental mortality. |
| Survival following release | $P(\text{surv} \mid \text{release})_{y,t} =$<br>$P(\text{release})_{y,t} \left(1 - \pi_{\text{FRIM}}\right)$ | Probability of survival following release. |
| Continuation probability | $P(\text{continue})_{y,t} = P(\text{none})_{y,t}$<br>$+ P(\text{surv} \mid \text{release})_{y,t}$ | Probability that a fish continues through the movement and natural-mortality process after either avoiding fishing or surviving release. The fishing outcomes satisfy $P(\text{continue}) + P(\text{FRIM}) + P(\text{kept}) = 1$. |

**Table S6:**
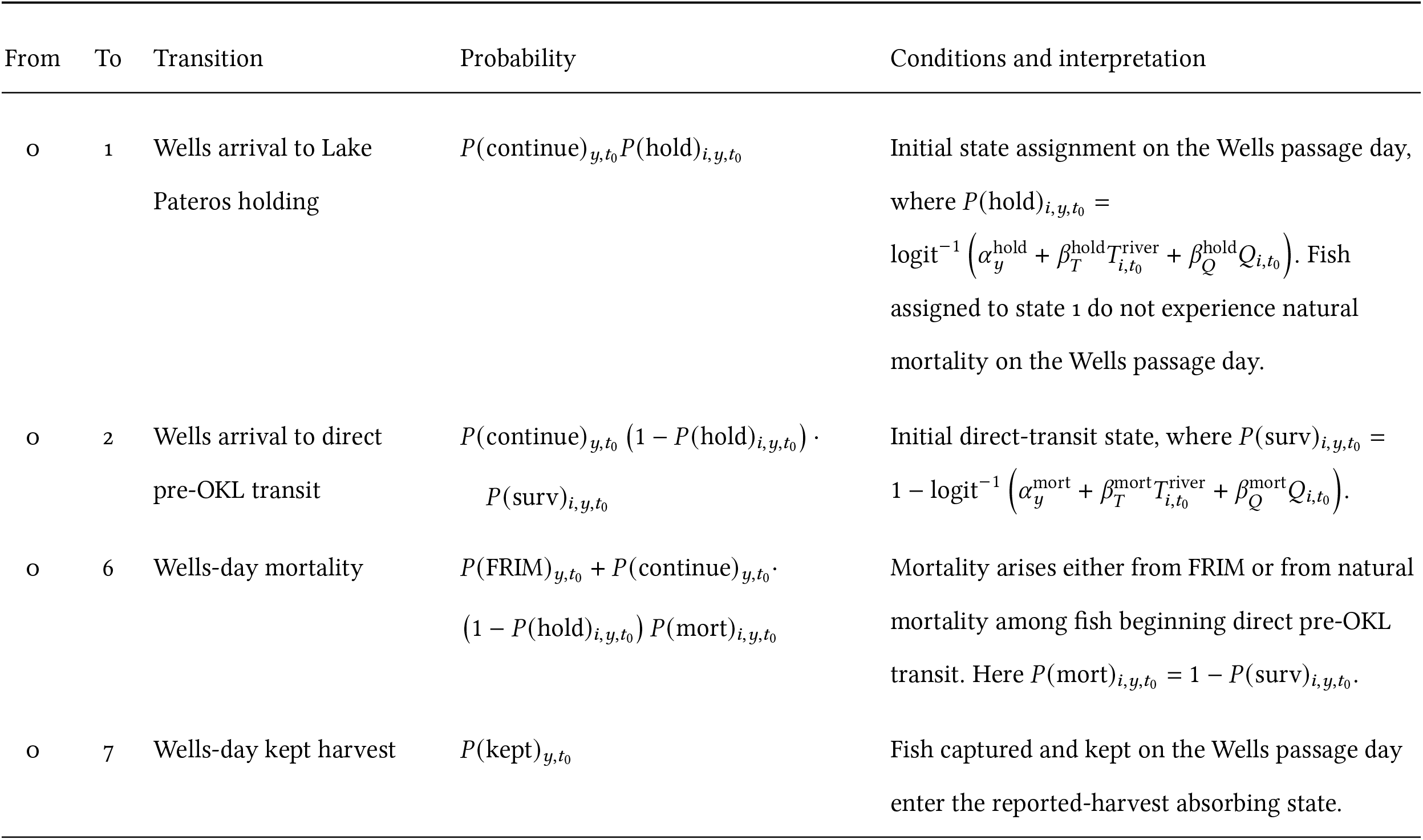
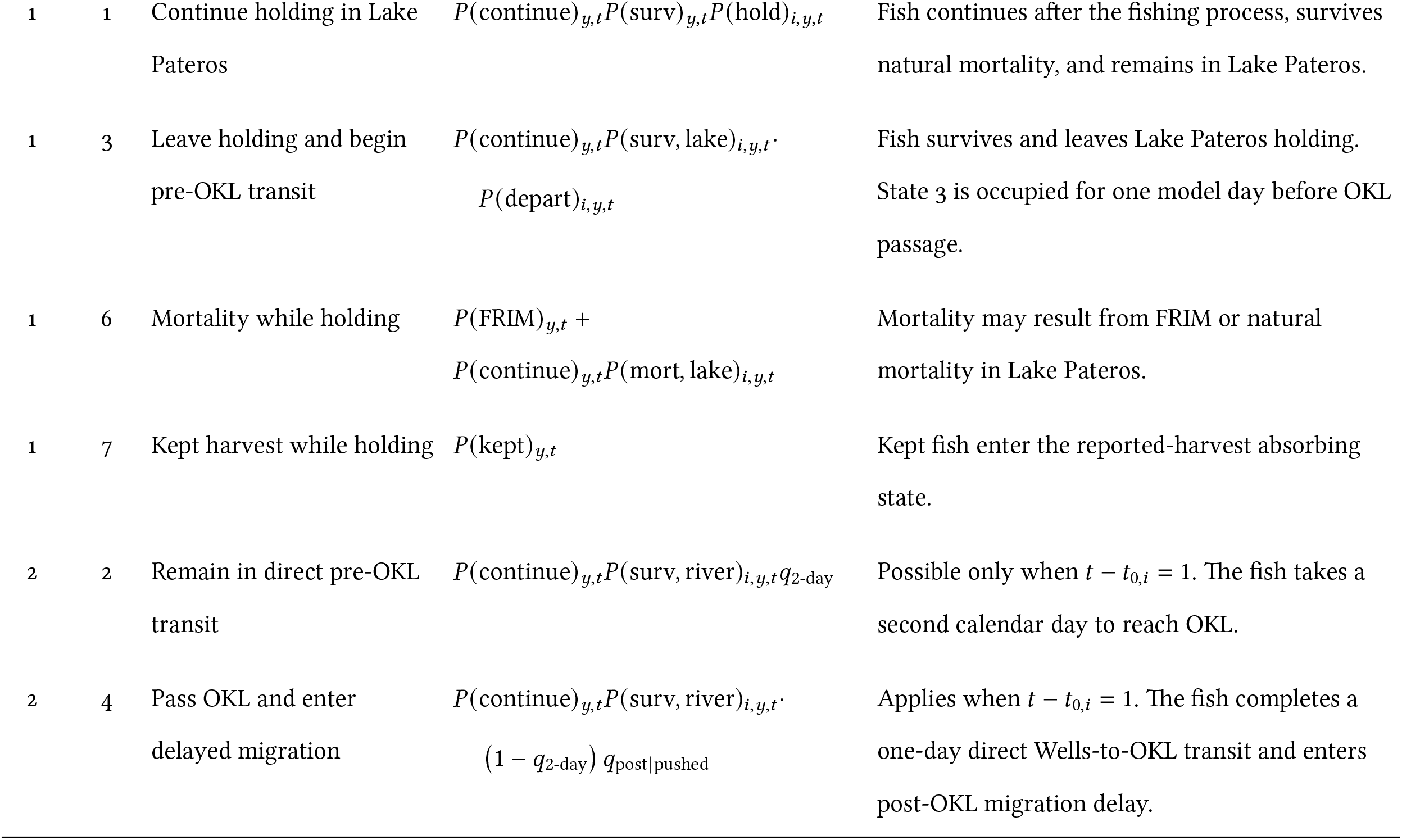
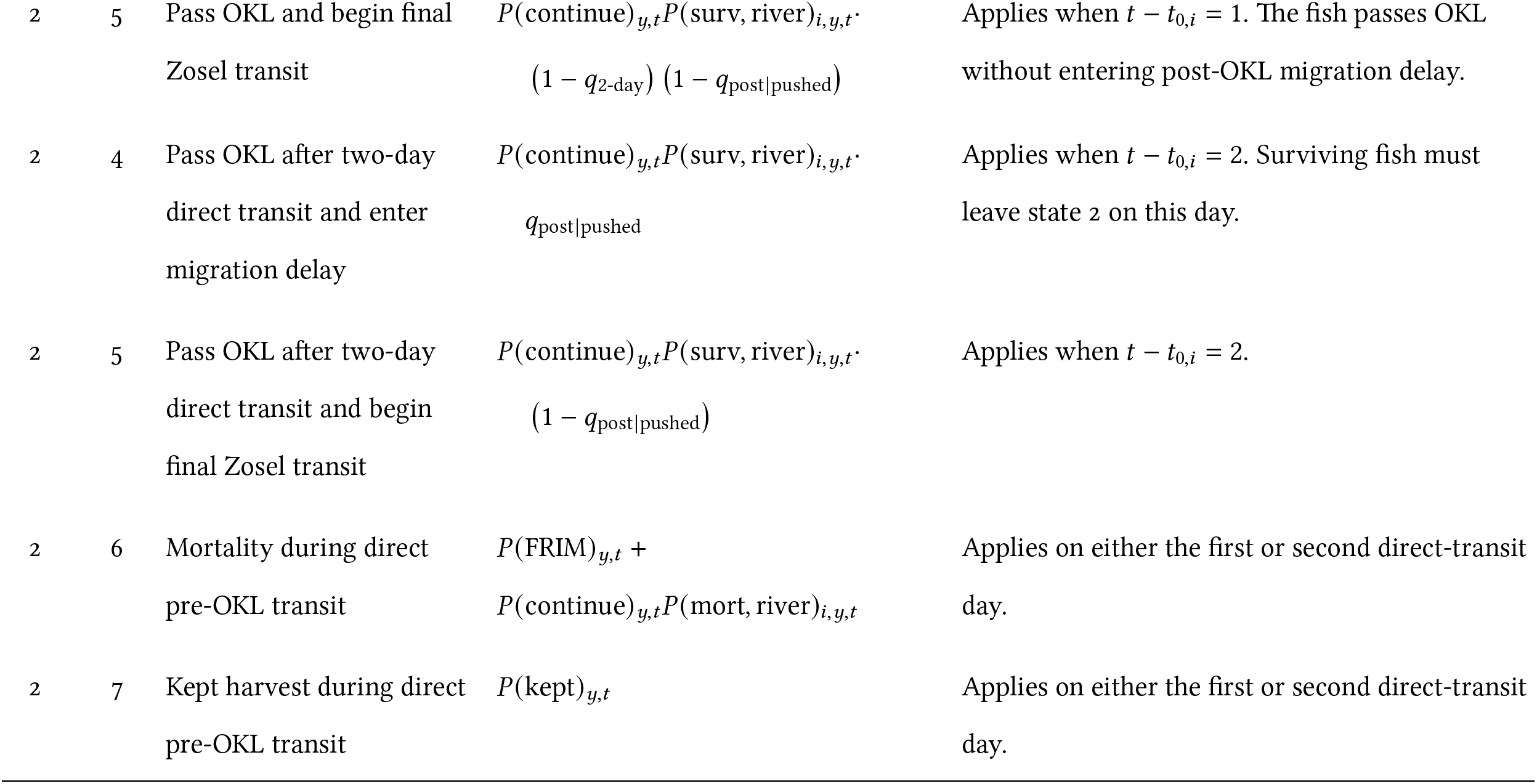
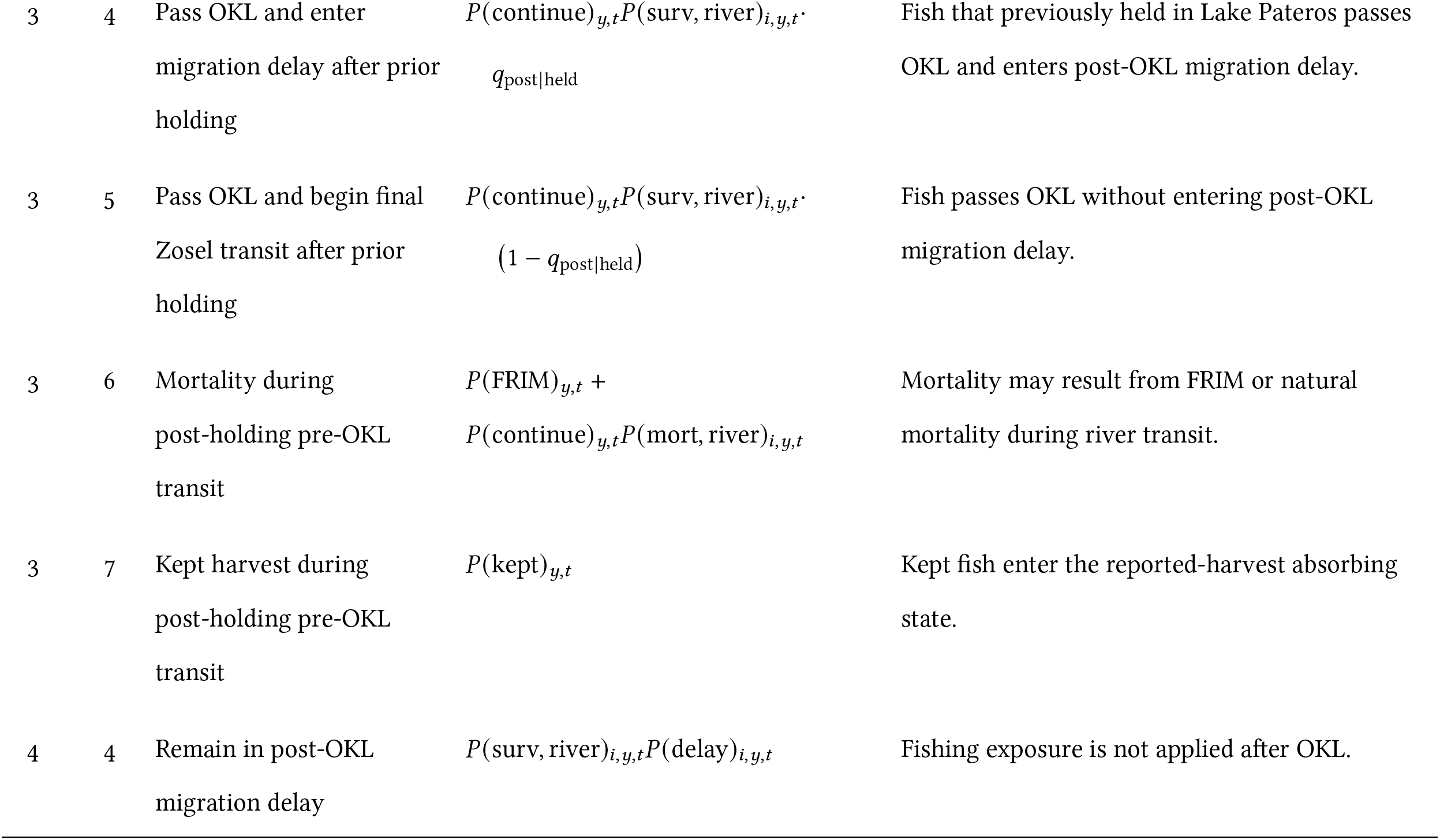
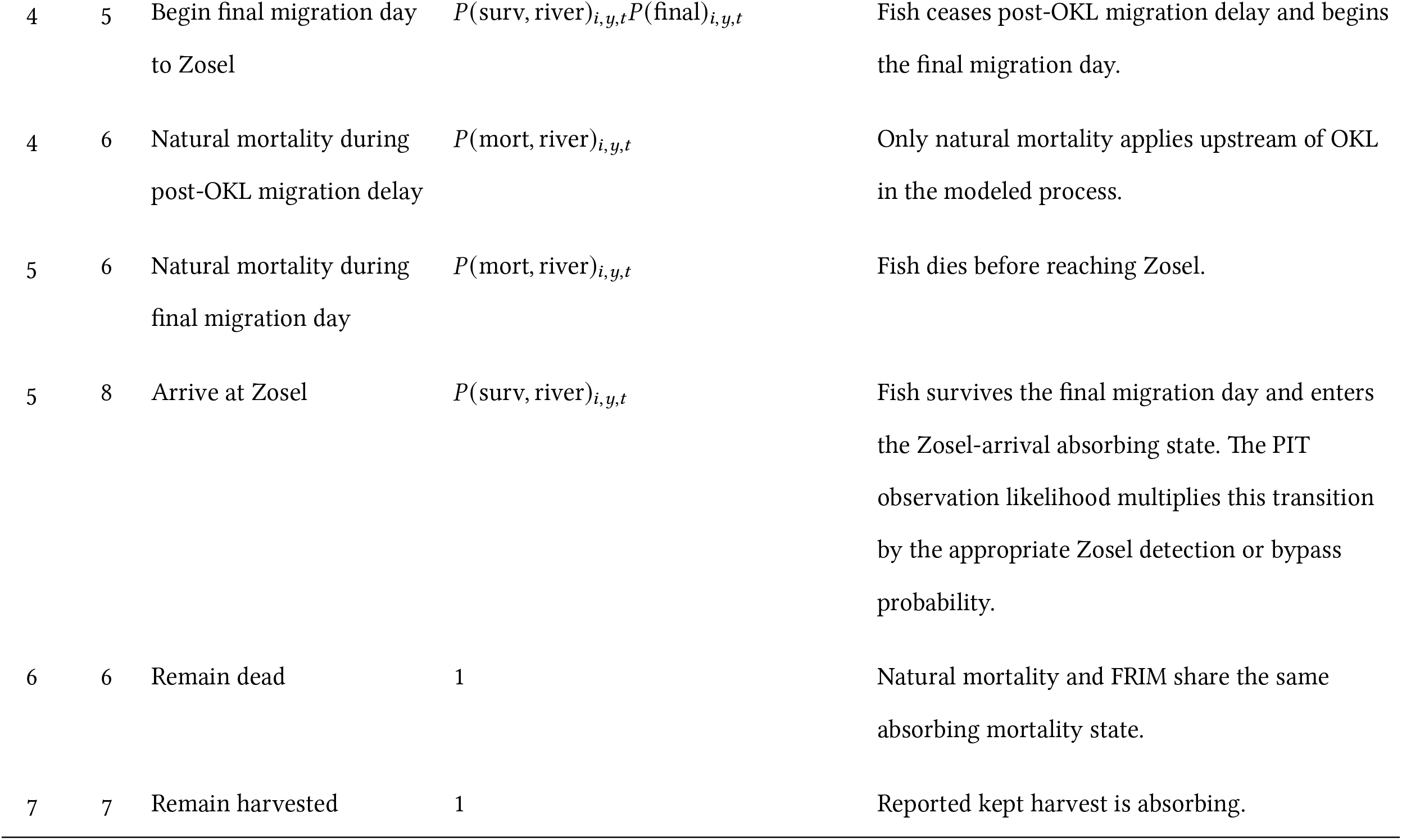
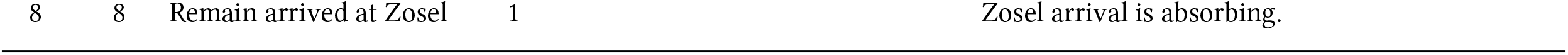
Daily latent-state transitions in the Wells-to-Zosel Sockeye migration model. Probabilities are shown before conditioning on PIT detections.

| From | To | Transition | Probability | Conditions and interpretation |
|---|---|---|---|---|
| 0 | 1 | Wells arrival to Lake Pateros holding | P(continue)<sub>y,t<sub>0</sub></sub>P(hold)<sub>i,y,t<sub>0</sub></sub> | Initial state assignment on the Wells passage day, where P(hold)<sub>i,y,t<sub>0</sub></sub> = logit<sup>−1</sup> (α<sub>y</sub><sup>hold</sup> + β<sub>T</sub><sup>hold</sup>T<sub>i,t<sub>0</sub></sub><sup>river</sup> + β<sub>Q</sub><sup>hold</sup>Q<sub>i,t<sub>0</sub></sub>). Fish assigned to state 1 do not experience natural mortality on the Wells passage day. |
| 0 | 2 | Wells arrival to direct pre-OKL transit | P(continue)<sub>y,t<sub>0</sub></sub> (1 − P(hold)<sub>i,y,t<sub>0</sub></sub>) · P(surv)<sub>i,y,t<sub>0</sub></sub> | Initial direct-transit state, where P(surv)<sub>i,y,t<sub>0</sub></sub> = 1 − logit<sup>−1</sup> (α<sub>y</sub><sup>mort</sup> + β<sub>T</sub><sup>mort</sup>T<sub>i,t<sub>0</sub></sub><sup>river</sup> + β<sub>Q</sub><sup>mort</sup>Q<sub>i,t<sub>0</sub></sub>). |
| 0 | 6 | Wells-day mortality | P(FRIM)<sub>y,t<sub>0</sub></sub> + P(continue)<sub>y,t<sub>0</sub></sub>· (1 − P(hold)<sub>i,y,t<sub>0</sub></sub>) P(mort)<sub>i,y,t<sub>0</sub></sub> | Mortality arises either from FRIM or from natural mortality among fish beginning direct pre-OKL transit. Here P(mort)<sub>i,y,t<sub>0</sub></sub> = 1 − P(surv)<sub>i,y,t<sub>0</sub></sub>. |
| 0 | 7 | Wells-day kept harvest | P(kept)<sub>y,t<sub>0</sub></sub> | Fish captured and kept on the Wells passage day enter the reported-harvest absorbing state. |
| 1 | 1 | Continue holding in Lake Pateros | P(continue)<sub>y,t</sub>P(surv)<sub>y,t</sub>P(hold)<sub>i,y,t</sub> | Fish continues after the fishing process, survives natural mortality, and remains in Lake Pateros. |
| 1 | 3 | Leave holding and begin pre-OKL transit | P(continue)<sub>y,t</sub>P(surv, lake)<sub>i,y,t</sub>· P(depart)<sub>i,y,t</sub> | Fish survives and leaves Lake Pateros holding. State 3 is occupied for one model day before OKL passage. |
| 1 | 6 | Mortality while holding | P(FRIM)<sub>y,t</sub> + P(continue)<sub>y,t</sub>P(mort, lake)<sub>i,y,t</sub> | Mortality may result from FRIM or natural mortality in Lake Pateros. |
| 1 | 7 | Kept harvest while holding | P(kept)<sub>y,t</sub> | Kept fish enter the reported-harvest absorbing state. |
| 2 | 2 | Remain in direct pre-OKL transit | P(continue)<sub>y,t</sub>P(surv, river)<sub>i,y,t</sub>q<sub>2-day</sub> | Possible only when t − t<sub>0,i</sub> = 1. The fish takes a second calendar day to reach OKL. |
| 2 | 4 | Pass OKL and enter delayed migration | P(continue)<sub>y,t</sub>P(surv, river)<sub>i,y,t</sub>· (1 − q<sub>2-day</sub>) q<sub>post\|pushed</sub> | Applies when t − t<sub>0,i</sub> = 1. The fish completes a one-day direct Wells-to-OKL transit and enters post-OKL migration delay. |
| 2 | 5 | Pass OKL and begin final Zosel transit | P(continue)<sub>y,t</sub>P(surv, river)<sub>i,y,t</sub>· (1 − q<sub>2-day</sub>) (1 − q<sub>post\|pushed</sub>) | Applies when t − t<sub>0,i</sub> = 1. The fish passes OKL without entering post-OKL migration delay. |
| 2 | 4 | Pass OKL after two-day direct transit and enter migration delay | P(continue)<sub>y,t</sub>P(surv, river)<sub>i,y,t</sub>· q<sub>post\|pushed</sub> | Applies when t − t<sub>0,i</sub> = 2. Surviving fish must leave state 2 on this day. |
| 2 | 5 | Pass OKL after two-day direct transit and begin final Zosel transit | P(continue)<sub>y,t</sub>P(surv, river)<sub>i,y,t</sub>· (1 − q<sub>post\|pushed</sub>) | Applies when t − t<sub>0,i</sub> = 2. |
| 2 | 6 | Mortality during direct pre-OKL transit | P(FRIM)<sub>y,t</sub> + P(continue)<sub>y,t</sub>P(mort, river)<sub>i,y,t</sub> | Applies on either the first or second direct-transit day. |
| 2 | 7 | Kept harvest during direct pre-OKL transit | P(kept)<sub>y,t</sub> | Applies on either the first or second direct-transit day. |
| 3 | 4 | Pass OKL and enter migration delay after prior holding | P(continue)<sub>y,t</sub>P(surv, river)<sub>i,y,t</sub>· q<sub>post\|held</sub> | Fish that previously held in Lake Pateros passes OKL and enters post-OKL migration delay. |
| 3 | 5 | Pass OKL and begin final Zosel transit after prior holding | P(continue)<sub>y,t</sub>P(surv, river)<sub>i,y,t</sub>· (1 − q<sub>post\|held</sub>) | Fish passes OKL without entering post-OKL migration delay. |
| 3 | 6 | Mortality during post-holding pre-OKL transit | P(FRIM)<sub>y,t</sub> + P(continue)<sub>y,t</sub>P(mort, river)<sub>i,y,t</sub> | Mortality may result from FRIM or natural mortality during river transit. |
| 3 | 7 | Kept harvest during post-holding pre-OKL transit | P(kept)<sub>y,t</sub> | Kept fish enter the reported-harvest absorbing state. |
| 4 | 4 | Remain in post-OKL migration delay | P(surv, river)<sub>i,y,t</sub>P(delay)<sub>i,y,t</sub> | Fishing exposure is not applied after OKL. |
| 4 | 5 | Begin final migration day to Zosel | P(surv, river)<sub>i,y,t</sub>P(final)<sub>i,y,t</sub> | Fish ceases post-OKL migration delay and begins the final migration day. |
| 4 | 6 | Natural mortality during post-OKL migration delay | P(mort, river)<sub>i,y,t</sub> | Only natural mortality applies upstream of OKL in the modeled process. |
| 5 | 6 | Natural mortality during final migration day | P(mort, river)<sub>i,y,t</sub> | Fish dies before reaching Zosel. |
| 5 | 8 | Arrive at Zosel | P(surv, river)<sub>i,y,t</sub> | Fish survives the final migration day and enters the Zosel-arrival absorbing state. The PIT observation likelihood multiplies this transition by the appropriate Zosel detection or bypass probability. |
| 6 | 6 | Remain dead | 1 | Natural mortality and FRIM share the same absorbing mortality state. |
| 7 | 7 | Remain harvested | 1 | Reported kept harvest is absorbing. |
| 8 | 8 | Remain arrived at Zosel | 1 | Zosel arrival is absorbing. |

**Table S7:** Observation constraints applied to the latent-state transitions.

| Observation | Likelihood contribution | State or transition constraint |
|---|---|---|
| OKL detected | Transitions 2 → 4, 2 → 5, 3 → 4, and 3 → 5 on the recorded OKL day are multiplied by *p*<sub>OKL,*y*</sub>. | OKL-crossing transitions are prohibited on all other days. Before the recorded OKL day, post-OKL and absorbing states are prohibited. On the OKL day, the fish must end in state 4 or 5. |
| OKL not detected while operating | Every permitted OKL-crossing transition is multiplied by 1 − *p*<sub>OKL,*y*</sub>. | The fish may cross OKL on any biologically permissible day. |
| OKL not operating | The OKL bypass contribution equals one. | No OKL detection information is applied. |
| Zosel detected | The transition 5 → 8 on the recorded Zosel day is multiplied by *p*<sub>ZSL,*y*</sub>. | The fish cannot arrive at Zosel on another day. Before the recorded Zosel day, mortality, harvest, and arrival are prohibited. On the preceding day, the fish must occupy state 5; on the detection day, the fish must enter state 8. |
| Zosel not detected while operating | Every permitted transition 5 → 8 is multiplied by 1 − *p*<sub>ZSL,*y*</sub>. | Undetected Zosel arrival may occur on any biologically permissible day. |
| Zosel not operating | The Zosel bypass contribution equals one. | No Zosel detection information is applied. |
| UOK detected | State 8 receives the terminal factor *ϕ*<sub>*y*</sub><sup>ZSL→UOK</sup>*p*<sub>UOK,*y*</sub>, where *ϕ*<sub>*y*</sub><sup>*ZSL*→*UOK*</sup> is survival between Zosel and the upstream array aggregate. | Only state 8 is compatible with a UOK detection; all other terminal states are prohibited. When Zosel was not detected, the latent process is propagated only through the recorded UOK day. |
| UOK not detected while operating | Terminal state 8 receives the factor (1 − *ϕ*<sub>*y*</sub><sup>ZSL→UOK</sup>) + *ϕ*<sub>*y*</sub><sup>ZSL→UOK</sup> (1 − *p*<sub>UOK,*y*</sub>) | A fish that reached Zosel may subsequently become unavailable before UOK or remain available but be missed. Terminal mortality and harvest states remain compatible with bypass. |
| UOK not operating | The UOK bypass contribution equals one. | No UOK detection information is applied. |
| Terminal-state restriction | No additional probability factor. | At the end of the represented trajectory, states 1–5 are assigned zero probability. Fish must occupy mortality, reported harvest, or Zosel arrival states. |

## References

Alexander, C.A.D., Alameddine, I., Machin, D., and Alex, K. 2024. A weight-of-evidence approach for understanding the recovery of Okanagan sockeye salmon. Environmental Management 74(6): 1063–1085. doi:10.1007/s00267-024-02031-y.

Bailey, C.J., Stiff, H.W., Thompson, P.L., Judson, B., and Ogden, A.D. 2025. Osoyoos Lake sockeye salmon (*Oncorhynchus nerka*) return abundance (1977–2023), adult age composition (1985– 2023), and marine survival (1998–2021). Canadian Manuscript Report of Fisheries and Aquatic Sciences 3282: vi +49p. 10.60825/v3g8-jw96.

Banks, J.W. 1969. A review of the literature on the upstream migration of adult salmonids. Journal of Fish Biology 1(2): 85–136. doi:10.1111/j.1095-8649.1969.tb03847.x.

Berdahl, A., Westley, P.A., Levin, S.A., Couzin, I.D., and Quinn, T.P. 2016. A collective navigation hypothesis for homeward migration in anadromous salmonids. Fish and Fisheries 17(2): 525–542. doi:10.1111/faf.12084.

Berdahl, A., Westley, P.A., and Quinn, T.P. 2017. Social interactions shape the timing of spawning migrations in an anadromous fish. Animal Behaviour 126: 221–229. doi:10.1016/j.anbehav.2017.01.020.

von Biela, V.R., Sergeant, C.J., Carey, M.P., Liller, Z., Russell, C., Quinn-Davidson, S., Rand, P.S., Westley, P.A., and Zimmerman, C.E. 2022. Premature mortality observations among Alaska’s Pacific salmon during record heat and drought in 2019. Fisheries 47(4): 157–168. doi:10.1002/fsh.10705.

Carpenter, B., Gelman, A., Hoffman, M.D., Lee, D., Goodrich, B., Betancourt, M., Brubaker, M., Guo, J., Li, P., and Riddell, A. 2017. *Stan* : A probabilistic programming language. Journal of Statistical Software 76(1). doi:10.18637/jss.v076.i01.

Crossin, G.T., Hinch, S.G., Farrell, A.P., Higgs, D.A., Lotto, A.G., Oakes, J.D., and Healey, M.C. 2004. Energetics and morphology of sockeye salmon: effects of upriver migratory distance and elevation. Journal of Fish Biology 65(3): 788–810. doi:10.1111/j.0022-1112.2004.00486.x.

Crozier, L.G., McClure, M.M., Beechie, T., Bograd, S.J., Boughton, D.A., Carr, M., Cooney, T.D., Dunham, J.B., Greene, C.M., Haltuch, M.A., Hazen, E.L., Holzer, D.M., Huff, D.D., Johnson, R.C., Jordan, C.E., Kaplan, I.C., Lindley, S.T., Mantua, N.J., Moyle, P.B., Myers, J.M., Nelson, M.W., Spence, B.C., Weitkamp, L.A., Williams, T.H., and Willis-Norton, E. 2019. Climate vulnerability assessment for Pacific salmon and steelhead in the California Current Large Marine Ecosystem. PLOS ONE 14(7): e0217711. doi:10.1371/journal.pone.0217711.

Crozier, L.G., Siegel, J.E., Wiesebron, L.E., Trujillo, E.M., Burke, B.J., Sandford, B.P., and Widener, D.L. 2020. Snake River sockeye and chinook salmon in a changing climate: Implications for upstream migration survival during recent extreme and future climates. PLOS ONE 15(9): e0238886. doi:10.1371/journal.pone.0238886.

Curasi, S.R., Melton, J.R., Arora, V.K., Humphreys, E.R., and Whaley, C.H. 2024. Global climate change below 2°C avoids large end century increases in burned area in Canada. npj Climate and Atmospheric Science 7(1). doi:10.1038/s41612-024-00781-4.

David, A.T., Asarian, J.E., and Lake, F.K. 2018. Wildfire smoke cools summer river and stream water temperatures. Water Resources Research 54(10): 7273–7290.

DeCicco, L., Hirsch, R., Lorenz, D., Watkins, D., Blodgett, D., Johnson, M., Zemmels, J., and Hinman, E. 2014. dataretrieval: Retrieval functions for USGS and EPA hydrology and water quality data. doi:10.32614/cran.package.dataretrieval.

Eliason, E.J., Clark, T.D., Hague, M.J., Hanson, L.M., Gallagher, Z.S., Jeffries, K.M., Gale, M.K., Patterson, D.A., Hinch, S.G., and Farrell, A.P. 2011. Differences in thermal tolerance among sockeye salmon populations. Science 332(6025): 109–112. doi:10.1126/science.1199158.

Farrell, A., Hinch, S., Cooke, S., Patterson, D., Crossin, G., Lapointe, M., and Mathes, M. 2008. Pacific salmon in hot water: Applying aerobic scope models and biotelemetry to predict the success of spawning migrations. Physiological and Biochemical Zoology 81(6): 697–709. doi: 10.1086/592057.

Fisheries and Oceans Canada 2025. Integrated Fisheries Management Plan for Salmon of Southern British Columbia: July 1, 2025 – June 30, 2026.

Fryer, J.K., Kelsey, D., Folks, S., Bussanich, R., Hyatt, K.D., and Stockwell, M.M. 2017. Studies into factors limiting the abundance of Okanagan and Wenatchee sockeye salmon in 2015. Columbia River Inter-Tribal Fish Commission Technical Report 17-06: xvi +199p.

Gabry, J., Češnovar, R., Johnson, A., and Bronder, S. 2025. cmdstanr: R Interface to ‘CmdStan’.

Gneiting, T. and Raftery, A.E. 2007. Strictly proper scoring rules, prediction, and estimation. Journal of the American Statistical Association 102(477): 359–378. doi:10.1198/016214506000001437.

Goniea, T.M., Keefer, M.L., Bjornn, T.C., Peery, C.A., Bennett, D.H., and Stuehrenberg, L.C. 2006. Behavioral thermoregulation and slowed migration by adult fall chinook salmon in response to high Columbia River water temperatures. Transactions of the American Fisheries Society 135(2): 408–419. doi:10.1577/t04-113.1.

Gustafson, R.G., Waples, R.S., Myers, J.M., Weitkamp, L.A., Bryant, G.J., Johnson, O.W., and Hard, J.J. 2007. Pacific salmon extinctions: Quantifying lost and remaining diversity. Conservation Biology 21(4): 1009–1020. doi:10.1111/j.1523-1739.2007.00693.x.

Hague, M.J., Ferrari, M.R., Miller, J.R., Patterson, D.A., Russell, G.L., Farrell, A.P., and Hinch, S.G. 2011. Modelling the future hydroclimatology of the lower Fraser River and its impacts on the spawning migration survival of sockeye salmon. Global Change Biology 17(1): 87–98. doi: 10.1111/j.1365-2486.2010.02225.x.

Holsman, K.K., Hazen, E.L., Haynie, A., Gourguet, S., Hollowed, A., Bograd, S.J., Samhouri, J.F., and Aydin, K. 2019. Towards climate resiliency in fisheries management. ICES Journal of Marine Science doi:10.1093/icesjms/fsz031.

Hyatt, K.D. and Rankin, D.P. 1999. A habitat based evaluation of Okanagan sockeye salmon escapement objectives. Canadian Science Advisory Secretariat Research Document 99/191: 59p.

Hyatt, K.D. and Stockwell, M.M. 2019. Chasing an Illusion? Successful Restoration of Okanagan River Sockeye Salmon in a Sea of Uncertainty. In From Catastrophe to Recovery: Stories of Fisheries Management Successes, edited by C.C. Krueger, W.W. Taylor, and S. Youn, American Fisheries Society, Bethesda, Maryland, pp. 65–100.

Hyatt, K.D., Stockwell, M.M., and Rankin, D.P. 2003. Impact and adaptation responses of okanagan river sockeye salmon (*Oncorhynchus nerka*) to climate variation and change effects during freshwater migration: Stock restoration and fisheries management implications. Canadian Water Resources Journal 28(4): 689–713. doi:10.4296/cwrj2804689.

Hyatt, K.D., Stiff, H.W., and Stockwell, M.M. 2020. Historic water temperature (1924–2018), river discharge (1929–2018), and adult sockeye salmon migration (1937–2018) observations in the Columbia, Okanogan, and Okanagan rivers. Canadian Manuscript Report of Fisheries and Aquatic Sciences 3206: xv +203p.

Macdonald, J.S., Patterson, D.A., Hague, M.J., and Guthrie, I.C. 2010. Modeling the influence of environmental factors on spawning migration mortality for sockeye salmon fisheries management in the Fraser River, British Columbia. Transactions of the American Fisheries Society 139(3): 768–782. doi:10.1577/t08-223.1.

Major, R.L. and Mighell, J.L. 1966. Influence of Rocky Reach Dam and the temperature of the Okanogan River on the upstream migration of sockeye salmon. Fishery Bulletin of the U.S. Fish and Wildlife Service 66: 131–147.

Marlier, M.E., Xiao, M., Engel, R., Livneh, B., Abatzoglou, J.T., and Lettenmaier, D.P. 2017. The 2015 drought in Washington State: a harbinger of things to come? Environmental Research Letters 12(11): 114008. doi:10.1088/1748-9326/aa8fde.

Martins, E.G., Hinch, S.G., Patterson, D.A., Hague, M.J., Cooke, S.J., Miller, K.M., LaPointe, M.F., English, K.K., and Farrell, A.P. 2011. Effects of river temperature and climate warming on stock-specific survival of adult migrating Fraser River sockeye salmon (*Oncorhynchus nerka*). Global Change Biology 17(1): 99–114. doi:10.1111/j.1365-2486.2010.02241.x.

Martins, E.G., Hinch, S.G., Cooke, S.J., and Patterson, D.A. 2012. Climate effects on growth, phenology, and survival of sockeye salmon (*Oncorhynchus nerka*): a synthesis of the current state of knowledge and future research directions. Reviews in Fish Biology and Fisheries 22(4): 887–914. doi:10.1007/s11160-012-9271-9.

Mayer, N.B., Hinch, S.G., and Eliason, E.J. 2024. Thermal tolerance in Pacific salmon: A systematic review of species, populations, life stages and methodologies. Fish and Fisheries 25(2): 283–302. doi:10.1111/faf.12808.

McClintock, B.T., Langrock, R., Gimenez, O., Cam, E., Borchers, D.L., Glennie, R., and Patterson, T.A. 2020. Uncovering ecological state dynamics with hidden Markov models. Ecology Letters 23(12): 1878–1903. doi:10.1111/ele.13610.

Murauskas, J., Hyatt, K.D., Fryer, J., Koontz, E., Folks, S., Bussanich, R., and Shelby, K. 2021. Migration and survival of Okanagan River sockeye salmon *Oncorhynchus nerka*, 2012–2019. Animal Biotelemetry 9(37): 1–16. doi:10.1186/s40317-021-00262-y.

Nass, B.L., Zapel, E., and Rowland, E. 2006. A conceptual proposal for a salmon enumeration facility on the Okanogan River. Technical report, LGL Limited.

Ogden, A., Alex, K., Pestal, G., Alameddine, I., Davis, B., Stiff, H.W., and Pham, S. 2025. Wild Salmon Policy Status, Limit Reference Point, and Candidate Escapement Goals for Okanagan Sockeye Salmon. Canadian Science Advisory Secretariat Research Document 2025/046: xii +102p.

Patterson, D.A., Robinson, K.A., Lennox, R.J., Nettles, T.L., Donaldson, L.A., Eliason, E.J., Raby, G.D., Chapman, J.M., Cook, K.V., Donaldson, M.R., Bass, A.L., Drenner, S.M., Reid, A.J., Cooke, S.J., and Hinch, S.G. 2017. Review and evaluation of fishing-related incidental mortality for Pacific salmon. Research Document 2017/010, DFO Can. Sci. Advis. Sec.

Polyakov, A.Y., Quinn, T.P., Myers, K.W., and Berdahl, A.M. 2022. Group size affects predation risk and foraging success in Pacific salmon at sea. Science Advances 8(26). doi:10.1126/sciadv.abm7548.

Pörtner, H.O. and Farrell, A.P. 2008. Physiology and climate change. Science 322(5902): 690–692. doi:10.1126/science.1163156.

Pörtner, H.O. and Knust, R. 2007. Climate change affects marine fishes through the oxygen limitation of thermal tolerance. Science 315(5808): 95–97. doi:10.1126/science.1135471.

Quinn, T.P., Hodgson, S., and Peven, C. 1997. Temperature, flow, and the migration of adult sockeye salmon (*Oncorhynchus nerka*) in the Columbia River. Canadian Journal of Fisheries and Aquatic Sciences 54(6): 1349–1360. doi:10.1139/f97-038.

R Core Team 2026. R: A Language and Environment for Statistical Computing. R Foundation for Statistical Computing. doi:10.32614/R.manuals.

Rand, P.S., Goslin, M., Gross, M.R., Irvine, J.R., Augerot, X., McHugh, P.A., and Bugaev, V.F. 2012. Global assessment of extinction risk to populations of sockeye salmon *Oncorhynchus nerka*. PLoS ONE 7(4): e34065. doi:10.1371/journal.pone.0034065.

Stiff, H.W. and Thompson, P.L. 2026. Projected water temperature and exposure indices for adult sockeye salmon migration in the Okanagan watershed, 2010–2033. Canadian Technical Report of Fisheries and Aquatic Sciences 3793: vii +102p. 10.60825/0jxx-3w53.

Thompson, P.L., Alex, K., Stiff, H.W., and Ogden, A.D. 2026. Temperature and flow impacts on spawner-to-fry production in Okanagan sockeye salmon. bioRxiv doi:10.64898/2026.09.25.754467.

van Vliet, M.T., Franssen, W.H., Yearsley, J.R., Ludwig, F., Haddeland, I., Lettenmaier, D.P., and Kabat, P. 2013. Global river discharge and water temperature under climate change. Global Environmental Change 23(2): 450–464. doi:10.1016/j.gloenvcha.2012.11.002.

Whitney, J.E., Al-Chokhachy, R., Bunnell, D.B., Caldwell, C.A., Cooke, S.J., Eliason, E.J., Rogers, M., Lynch, A.J., and Paukert, C.P. 2016. Physiological basis of climate change impacts on North American inland fishes. Fisheries 41(7): 332–345. doi:10.1080/03632415.2016.1186656.

Whoriskey, K., Martins, E.G., Auger-Méthé, M., Gutowsky, L.F.G., Lennox, R.J., Cooke, S.J., Power, M., and Mills Flemming, J. 2019. Current and emerging statistical techniques for aquatic telemetry data: A guide to analysing spatially discrete animal detections. Methods in Ecology and Evolution 10(7): 935–948. doi:10.1111/2041-210x.13188.

Williams, I.V. and Brett, J.R. 1987. Critical swimming speed of Fraser and Thompson River pink salmon (*Oncorhynchus gorbuscha*). Canadian Journal of Fisheries and Aquatic Sciences 44(2): 348–356. doi:10.1139/f87-043.

Wright, M.J., Hurson, M., Robinson, K.A., Patterson, D.A., and Venditti, J.G. 2025. A typology of potential hydraulic barriers to adult salmon migration in a bedrock river. Canadian Journal of Fisheries and Aquatic Sciences 82: 1–19. doi:10.1139/cjfas-2024-0100.

